# The Cortical Temporal Axis: MEG-Based Cross-Frequency Gradients with Biological Anchors

**DOI:** 10.1101/2025.11.07.687284

**Authors:** Xiaobo Liu, Xinyu Wu, Siyu Long, Lindsey Power, Dominic Boutet, Lang Liu, Zhenqi Liu, Ruiyang Ge, Sanwang Wang, Jianfeng Zhang, Li Dong, Tianyi Yan, Guoyuan Yang

## Abstract

Neural oscillations have long been used to characterize the temporal dynamics of individual brain regions, yet a parsimonious, system-level representation that integrates multi-rhythmic activity has been lacking. Using source-localized resting-state magnetoencephalography (MEG) data, we computed regional power spectra and assessed spectral similarity, then applied diffusion map embedding to construct cross-frequency neurophysiological gradients that place whole-brain oscillations within a unified, low-dimensional coordinate system. We found the first three gradients accounted for over 40% of the variance, remained stable across individuals, and aligned with established functional, structural, and geometric cortical axes. Computational modeling showed that these gradients reflect local excitation–inhibition balance, while multimodal analyses revealed strong associations with neurotransmitter receptor distributions, cytoarchitecture, and cell-type–specific gene expression. Lifespan analyses further demonstrated systematic gradient reorganization, with distinct cognitive mappings onto functions such as language, memory, and multisensory integration. Clinically, Parkinson’s disease patients displayed disrupted gradients, particularly in regions linked to language and social cognition. Finally, these gradient patterns exhibited high test–retest reliability. These findings establish MEG-derived neurophysiological gradients as robust, low-dimensional representations of cortical organization, offering a biologically grounded framework for studying brain aging and disease.

## Introduction

Neural oscillations are a fundamental property of brain dynamics, thought to play a central role in coordinating neuronal activity and supporting information integration (Zhang and Chen 2025). Distinct cortical regions exhibit characteristic dominant frequency patterns, such as α oscillations (7–13 Hz) in occipito-parietal areas and β oscillations (15–30 Hz) in the sensorimotor cortex (Hillebrand et al., 2016; Keitel & Gross, 2016). In contrast, δ (1–3 Hz) and θ (3–7 Hz) rhythms are more broadly distributed, particularly in frontal regions (Mellem et al., 2017). These region-specific rhythmic dynamics indicate that oscillations are not random noise but reflect systematic organizational principles of the cortex (van Es et al. 2025). Conventional time-series analyses in neuroscience have generally concentrated on power spectra under canonical electrophysiological rhythms. Recent advances in systems neuroscience have given rise to gradient-based methods for abstract representations of brain function, which capture functional dynamics across the cortex (van Es et al. 2025; Godfrey and Singh 2021; Mostame and Sadaghiani 2020). However, it remains unclear whether these multiple rhythmic features are embedded within the hierarchical architecture of the cortex, analogous to the macroscale gradients widely reported in anatomical and functional studies. Establishing whether oscillatory activity follows hierarchical spatial distributions would provide critical insights into how temporal dynamics support perception, memory, and cognitive control across distributed brain systems.

Accumulating evidence shows that the human cortex exhibits continuous gradients across multiple levels—cellular, anatomical, geometric, molecular, and functional gradients (Kim et al. 2024; Hansen and Misic 2025a; Hansen et al. 2021; Shafiei et al. 2023; Margulies et al. 2016a; Thomas Yeo et al. 2011). Along the posterior–anterior axis, neuronal density (Beul and Hilgetag 2019; Cahalane et al. 2012), cortical thickness (Valk et al. 2020), and myelination (Burt et al. 2018; Paquola et al. 2019) vary systematically, reflecting a transition from lower-order sensory regions dominated by feedforward projections to higher-order association areas characterized by feedback connectivity. Substantial evidence suggests that the dynamics of neural systems emerge from the structural excitations of the brain, which are inherently shaped by its multi-level microarchitecture (Paquola et al. 2020; Kong et al. 2021). The neural dynamics are also constrained by global geometry of the cortex, much like the shape of a drum or the length of a violin string determines its resonant frequencies (Pang et al. 2023). Crucially, this gradient organization extends beyond structure into temporal dynamics with sensory regions dominated by fast, high-frequency activity, whereas prefrontal areas engage slower rhythms that support cross-temporal integration (Murray et al. 2014). Neural oscillations therefore likely represent a critical “temporal axis” of cortical gradient organization. Their frequency distributions align with anatomical hierarchies and are further shaped by geometric constraints and molecular mechanisms, thereby bridging spatial gradients with functional stratification and serving as a key dynamical substrate linking brain structure and cognition (Shafiei et al. 2023; Hansen et al. 2022a). Previous studies have primarily used functional magnetic resonance imaging (fMRI) to establish spatiotemporal brain dynamics, which is temporally limited by the latency of the hemodynamic response (approximately 1s) (Belliveau et al. 1991; Kwong et al. 1992). Electroencephalography (EEG) or magnetoencephalography (MEG), on the other hand, directly measures the magnetic fields generated by neuronal activity and can provide temporal resolution of less than 1 ms, adequate for detecting the orchestration of complex cognitive activity (Shafiei et al. 2022). Therefore, utilizing resting state EEG/MEG spatiotemporal signals to conduct parsimonious characterization of neural oscillations will more effectively reflect key functional processes including sensory integration, attention, and memory (Hirano and Uhlhaas 2021; Beste et al. 2023).

Neural dynamic features also demonstrate systematic spatial structural variations across the human lifespan. For example, functional spectral profiles across different frequency bands undergo substantial reorganization from childhood to adolescence (Meng and Xiang 2016). In contrast, aging is characterized by changes in neural oscillations which vary as a function of brain region and frequency band (Park et al. 2025). The parsimonious macroscale functional architecture of the brain is typically inferred from haemodynamic activity (Margulies et al. 2016a). During adolescence, this macroscale representation undergoes a spatial developmental inversion driven by the ventral attention network, which reflects individual variability in cognitive ability (Y. Xia et al. 2022; Dong et al. 2024; Margulies et al. 2016b). In aging, this spatial configuration displays increasing dispersion of the frontoparietal, attention, and default mode networks—patterns that are negatively associated with cognitive performance, particularly fluid intelligence (Bethlehem et al. 2020). These observations indicate that the interpretation of ongoing neural activity is further complicated by its extensive, spatiotemporally heterogeneous associations with behavioral and physiological variables. Moreover, disruptions in the macroscale hierarchical organization—affecting the integration and segregation of unimodal and transmodal networks—can precipitate neurodevelopmental psychiatric conditions such as autism spectrum disorder (Hong et al. 2019) and depression (M. Xia et al. 2022). Similarly, in neurodegenerative diseases such as Parkinson’s disease (PD), the brain’s functional gradients exhibit atypical hierarchical organization coupled with distinct gene expression profiles (Wu et al. 2024; Li et al. 2025). Consequently, constructing parsimonious representations of neural oscillatory dynamics based on resting-state EEG/MEG holds substantial promise for elucidating the lifespan trajectories of brain cognitive functions, as well as for advancing our understanding of the neural mechanisms underlying neuropsychiatric and neurodegenerative disorders.

Here, we derived neurophysiological gradients from resting-state MEG data and examined their biological, functional, and clinical relevance. These gradients proved robust and aligned with functional, structural, and geometric axes, while also reflecting local excitation–inhibition balance and multi-scale features including receptor distributions, cytoarchitecture, and gene expression. Across the lifespan, gradients reorganized systematically with age and mapped onto distinct cognitive domains, with the first principal gradient showing the highest test–retest reliability. Clinically, Parkinson’s disease patients exhibited pronounced alterations in these gradients, particularly in networks supporting language, memory, and multisensory integration. Together, these findings establish MEG-derived gradients as low-dimensional representations of cortical organization and highlight their potential as biomarkers for brain aging and disease.

## Results

### Neurophysiological Gradient Construction and Basic Properties

The exploration cohort was Cam-CAN: 608 healthy adults (307 M/301 F; mean age 54.19 ± 18.19 years) with resting-state MEG and MRI. We first constructed neurophysiological gradients by computing source-level activity, estimating power spectra, and applying cosine similarity between cortical regions parcellated by the Schaefer 200 atlas, followed by diffusion map embedding (Fig. 1a).

**Fig. 1.**
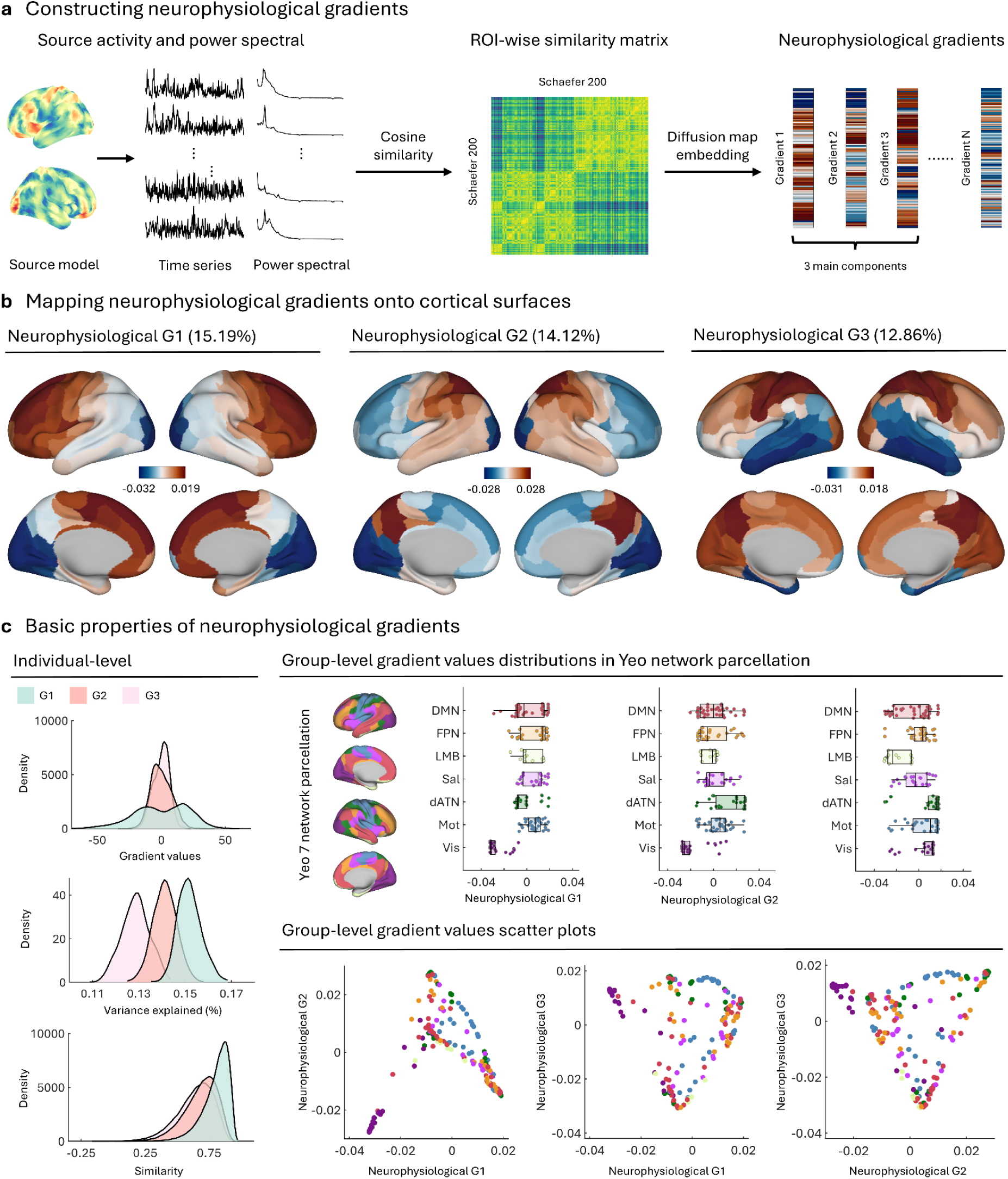
Construction and Basic Properties of Neurophysiological Gradients. **a.** Neurophysiological gradients were derived by computing source-level activity, estimating power spectra, calculating cosine similarity between regions (Schaefer 200 parcellation), and applying diffusion map embedding to extract gradient components. **b.** The first three gradients (G1–G3) are visualized on cortical surfaces, explaining 15.19%, 14.12%, and 12.86% of the variance, respectively. **c.** Gradient value distributions at the individual level, explained variance, similarity metrics, and group-level projections onto Yeo networks are shown. Scatter plots illustrate relationships among the three gradients.

The first three neurophysiological gradients (G1–G3) explained 15.19%, 14.12%, and 12.86% of the variance, respectively, and were projected onto cortical surfaces (Fig. 1b). Noted that higher-order components explain only limited additional variance and exhibit reduced stability and biological interpretability. G1 recapitulates the canonical primary sensory → association axis (low in visual/sensorimotor, high in DMN), G2 dissociates attentional/control systems from limbic–temporal territories, and G3 captures finer anterior–posterior and medial–lateral distinctions. Furthermore, we verified that these physiological gradients are not attributable to the source localization model (see **Supplementary Fig. 1**). At the individual level, gradient value distributions, explained variance, and similarity metrics indicated stable gradient representation (Fig. 1c, left). Group-level analysis revealed systematic projection of gradients onto canonical Yeo’s networks, with scatter plots further illustrating distinct interrelationships among first three neurophysiological gradients (Fig. 1c, right). Specifically, In G1, network values increase from Vis and Mot through Sal/LMB and dATN to FPN, with DMN highest, reflecting the canonical sensory–association axis. In G2, LMB is lowest and FPN/dATN highest, with DMN slightly negative, Vis/Mot near zero, and Sal slightly positive, indicating a control/attention versus limbic opposition. In G3, inter-network separation diminishes but distributions broaden, capturing fine-grained anterior–posterior and medial–lateral variation, with opposing orientations within DMN and FPN. Scatter plots further show a progression along G1 from Vis/Mot toward DMN/FPN, a pronounced separation of FPN/dATN from LMB along G2, and, on this basis, a more refined within-network differentiation by G3.

### Association Between Neurophysiological and Multimodal Cortical Gradients

To assess the relationship between neurophysiological gradients and established cortical organizational axes, we compared the first three neurophysiological gradients with multimodal gradients derived from fMRI, MRI and DTI, cortical geometry, and structural connectivity (Fig. 2a). Neurophysiological G1 showed a strong negative association with the first geometry mode (*r□* = −0.88) and a strong positive association with the first structural gradient (*r□* = 0.87). Neurophysiological G2 correlated positively with structure G2 (*r□* = 0.64) and showed a weak negative correlation with structure G3 (*r□* = −0.14). Neurophysiological G3 correlated negatively with function G1 (*r□* = −0.69) and with geometry mode2 (*r□* = −0.75). All reported effects survived FDR correction (*p* < 0.05).

**Fig. 2.**
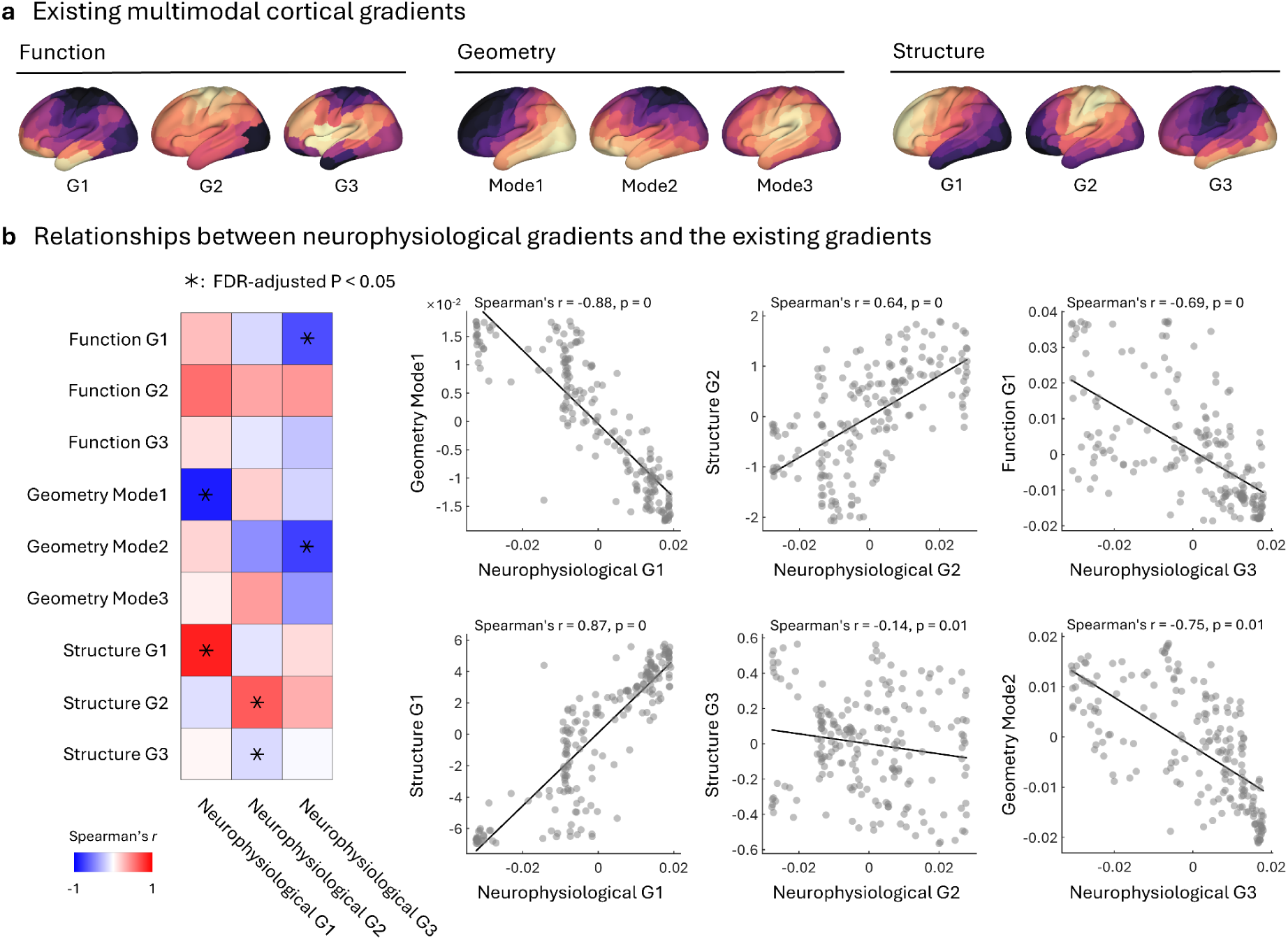
Correspondence Between Neurophysiological and Multimodal Cortical Gradients. **a.** Parsimonious cortical gradients derived from functional connectivity, geometry, and structure are displayed across three gradient modes of CAM-CAN dataset. Specifically, functional gradients were derived from resting-state functional connectivity (Function G1–G3), structural gradients from diffusion MRI–based structural connectivity (Structure G1–G3), and geometric modes correspond to the Laplace–Beltrami eigenfunctions of the cortical surface (Geometry Mode1–Mode3). **b.** Correlations between neurophysiological and multimodal gradients. The left heatmap and right scatter plots show Spearman correlations across the 200 cortical parcels. Statistical significance was assessed with a spin permutation test (10,000 rotations) to account for spatial autocorrelation and controlled using Benjamini–Hochberg FDR; significant effects are marked with asterisks (*FDR*-adjusted *p* < 0.05). Main results: — G1 was strongly negatively correlated with Geometry Mode1 (*r□* = −0.88) and strongly positively correlated with Structure G1 (*r□* = 0.87); — G2 correlated positively with Structure G2 (*r□* = 0.64) and weakly negatively with Structure G3 (*r□* = −0.14); — G3 showed negative correlations with Function G1 (*r□* = −0.69) and Geometry Mode2 (*r□* = −0.75).

Collectively, the results show that neurophysiological G1 mirrors the cortex’s principal structural/geomorphological axis, neurophysiological G2 indexes higher-order structural organization consistent with a shift toward control/association networks, and neurophysiological G3 aligns with the dominant functional axis and a secondary geometric mode, capturing complementary anterior–posterior and medial–lateral distinctions. Thus, MEG-derived neurophysiological gradients may capture additional, modality-specific information not fully reflected in structural or geometric axes, while still exhibiting clear correspondence with canonical macroscale cortical organization.

### Linking Neurophysiological Gradients to Simulated Brain Activity and E/I Balance

Here we asked whether the gradients index local circuit dynamics using a whole-brain Wilson–Cowan neural-mass model. Model-derived center frequencies reproduced the empirical MEG topography, showing a significant parcel-wise association at the group level (Fig.3.a Spearman r = 0.36, *p* = 0.01) and at the individual level (see bottom right of Fig.3.a). We then related the gradients to simulated excitation–inhibition (E/I) ratios which defined from average E and I activity within the simulated time series: only G2 showed a robust spatial coupling, exhibiting a negative correlation with E/I (r = −0.24, *p* = 0.02; results survive spin tests for spatial autocorrelation and FDR correction), whereas G1 and G3 were not significant. This pattern indicates that variation along G2—from primary toward control/association territories—tracks a shift toward lower E/I (relatively stronger inhibition), consistent with theoretical accounts linking E–I balance to oscillatory frequency and large-scale cortical dynamics

**Fig. 3.**
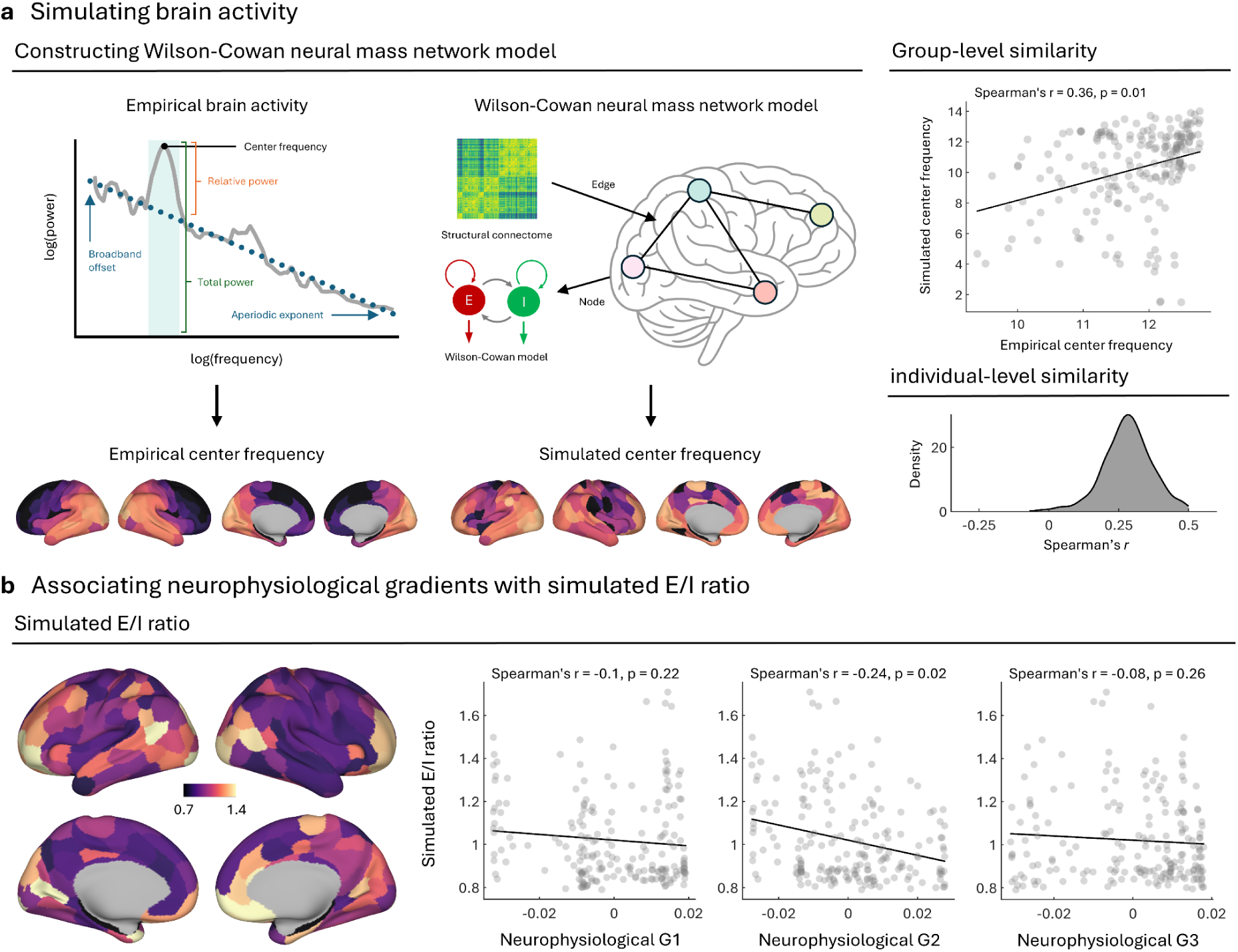
Linking Neurophysiological Gradients to Simulated Brain Activity and E/I Balance. **a.** Construction of a whole-brain network model based on Wilson–Cowan dynamics using structural connectome input. The simulated center frequency shows group-level correlation with empirical estimates (*r* = 0.36, *p* = 0.01). **b.** Spatial correlation between neurophysiological gradients and simulated excitation/inhibition (E/I) ratio maps. Neurophysiological G2 shows a modest but significant negative correlation with E/I ratio (*r* = –0.24, *p* = 0.02).

### Associations of Neurophysiological Gradients with Chemoarchitecture, Cytoarchitecture, and Cell Types

We examined the neurochemical and cytoarchitectonic bases of the MEG-derived gradients using PET receptor/transporter maps, BigBrain/quantitative-histology gradients, laminar thickness and cortical intensity profiles, and cell-type transcriptomic maps, all projected to Schaefer-200 parcels. Parcel-wise associations were quantified with Spearman correlations; significance was assessed with spin permutation (10,000 rotations) to account for spatial autocorrelation with FDR. Neurophysiological G1 showed significant associations with serotonergic (5-HT) and GABAergic receptor distributions, whereas G2 and G3 additionally related to dopaminergic and glutamatergic systems (Fig. 4a; FDR-adjusted *p* < 0.05). Cytoarchitecturally, G1 was strongly negatively correlated with the principal histological gradient (Fig. 4b, left; *r* = −0.81, *p* = 0), and G3 was positively correlated with the principal microstructural gradient (Fig. 4b, right; *r* = 0.60, *p* = 0). The gradients also mapped systematically onto laminar thickness (with deeper-layer effects for G1) and intensity profiles (Fig. 4b, bottom). Finally, cell-type transcriptomic analyses revealed selective associations with specific neuronal and glial populations—e.g., a link between G2 and deep-layer IT-type excitatory neurons—indicating that neurophysiological gradients reflect multi-scale cellular and molecular features of cortical organization (Fig. 4c).

**Fig. 4.**
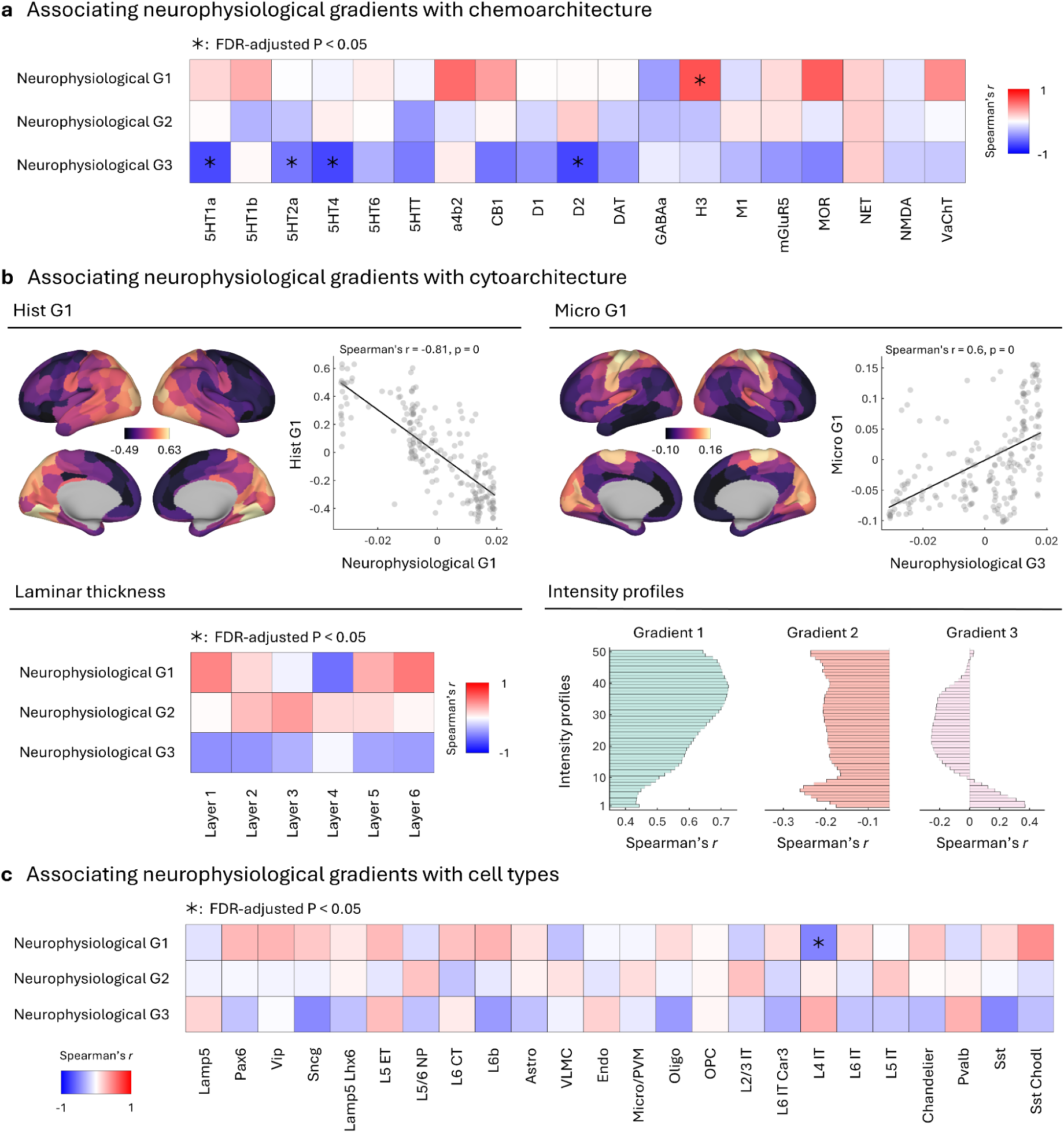
Associations of Neurophysiological Gradients with Chemoarchitecture, Cytoarchitecture, and Cell Types. **a.** Correlation heatmap showing the associations between gradients (G1–G3) and the spatial distribution of neurotransmitter receptor expression. Asterisks indicate statistically significant associations (FDR-adjusted *p* < 0.05). **b.** Associations with cytoarchitectonic features: neurophysiological G1 negatively correlates with the histological gradient (*r* = –0.81, *p* = 0); neurophysiological G3 positively correlates with the microstructural gradient (*r* = 0.6, *p* = 0). Relationships with cortical laminar thickness and intensity profiles are also presented. **c.** Correlation heatmap between neurophysiological gradients and gene expression profiles of specific cell types. Significant associations (FDR-adjusted *p* < 0.05) are marked with asterisks.

### Alterations of Neurophysiological Gradients in Aging and Parkinson’s Disease

Using the Cam-CAN dataset, we identified lifespan-wide associations between neurophysiological gradients and aging. Spatial maps revealed both positive and negative age-related effects, with red indicating increased gradient strength with age and blue indicating decreases (Fig. 5a). Cognitive term associations demonstrated that neurophysiological G1 aging effects were linked to language, action, and motor processes; neurophysiological G2 effects were associated with working memory, semantic memory, and social cognition; and neurophysiological G3 effects were linked to multisensory integration and attentional control (Fig. 5b). These findings suggest that age-related reorganization of neurophysiological gradients maps onto distinct cognitive domains.

**Fig. 5.**
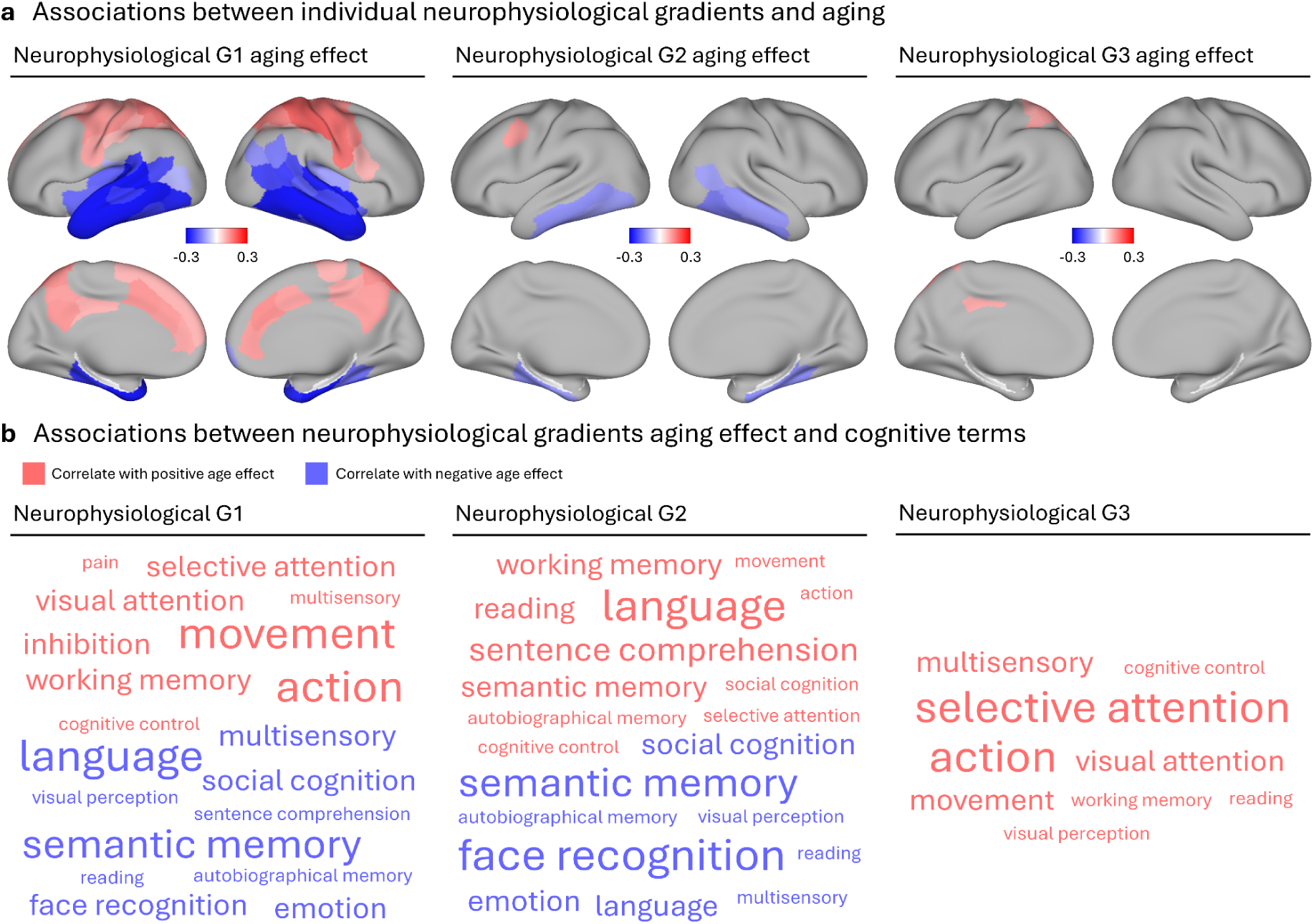
Associations Between Neurophysiological Gradients, Aging, and Cognitive Terms. **a.** Cortical projections showing associations between each neurophysiological gradient (G1–G3) and age. Red indicates a positive age effect (increase with age), and blue indicates a negative age effect (decrease with age). **b.** Word clouds representing cognitive functions whose spatial maps correlate with the age-related effects of each gradient. Red terms indicate positive age associations; blue terms indicate negative age associations.

Furthermore, we compared neurophysiological gradients between PD patients and healthy controls (Supplementary Fig. 4). Significant differences were observed in neurophysiological G1–G3, with PD patients exhibiting both increased and decreased gradient values in distinct cortical regions (Supplementary Fig. 5a). Cognitive associations revealed that alterations in neurophysiological G1 was linked to reading, semantic memory, pain, and language; neurophysiological G2 alterations corresponded to social cognition, autobiographical memory, and emotion; while neurophysiological G3 alterations mapped onto multisensory integration, attention, and action control (Supplementary Fig. 5b). These results indicate that PD is characterized by widespread disruptions in neurophysiological gradients linked to higher-order cognitive dysfunction.

### Test–Retest Reliability and Reproducibility of Neurophysiological Gradients

This study utilized data from the OMEGA platform comprising 158 unrelated participants (77 females; mean age = 31.9 ± 14.7 years), each contributing approximately 5 minutes of recording; a subset of 47 individuals underwent multiple sessions on separate days. We quantified test–retest reliability from repeated within-subject MEG recordings using intraclass correlation coefficients (ICC) computed on parcel-wise gradient scores after running the identical preprocessing and source/gradient pipeline for each session (Fig. 6a).

**Fig. 6.**
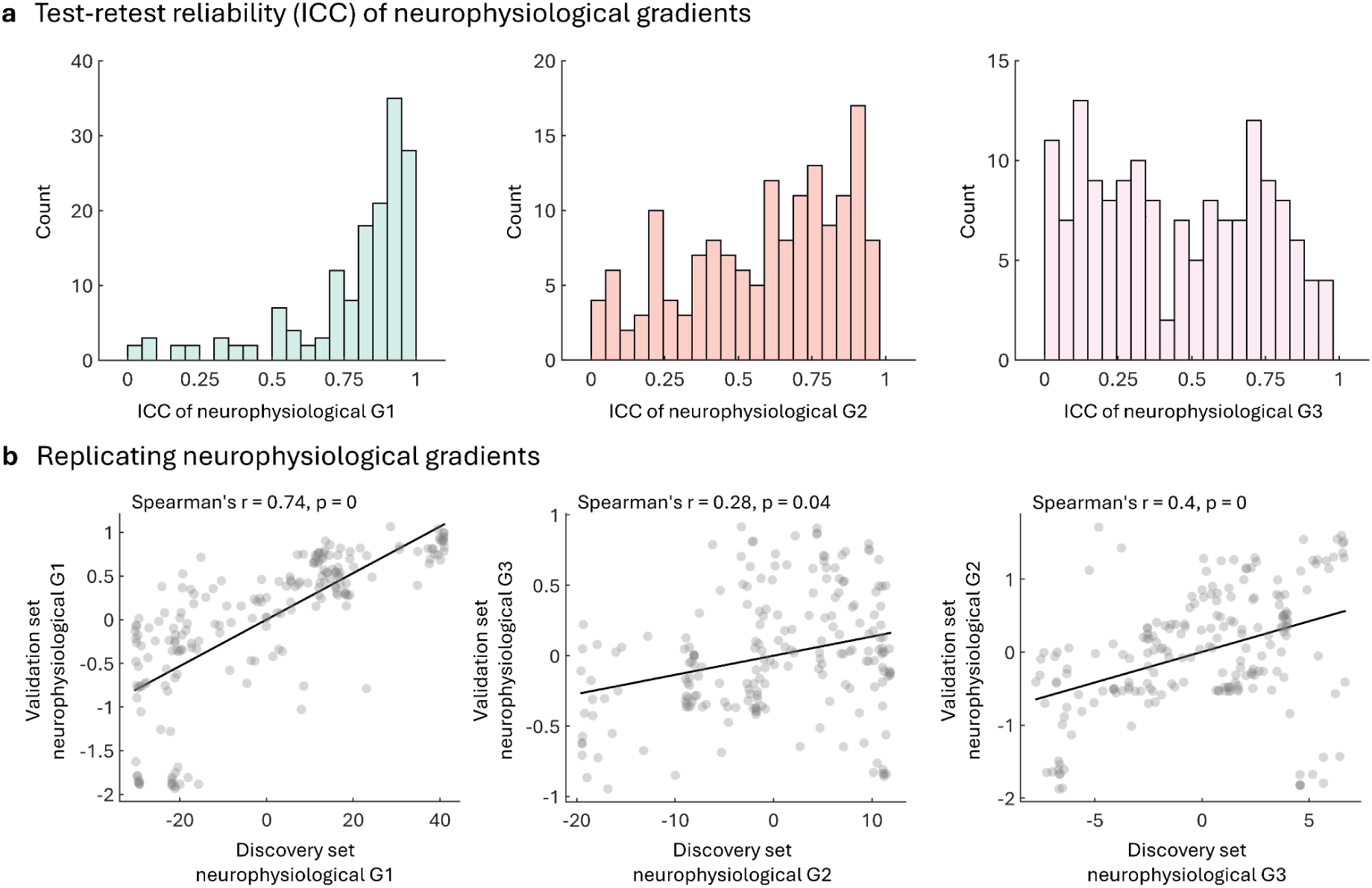
Test-Retest Reliability and Reproducibility of Neurophysiological Gradients. **a.** Distribution of intraclass correlation coefficients (ICCs) across individuals for neurophysiological G1–G3, with G1 shows the highest test-retest reliability. **b.** Replication of neurophysiological gradients in an independent validation dataset. Spearman’s rank correlations between discovery and validation sets are highest for neurophysiological G1 (*r* = 0.74, *p* = 0), and moderate for neurophysiological G2 (*r* = 0.28, *p* = 0.05) and G3 (ρ = 0.4, *p* = 0).

G1 showed the highest stability, with ICCs concentrated near 1, whereas G2 and G3 displayed broader, moderate ICC distributions. For external replication, we used an independent validation cohort processed with the same steps (source reconstruction → PSD → spectrum-based similarity → diffusion embedding). Group-level gradients were derived separately in the discovery and validation datasets; to remove sign/axis ambiguity, embeddings were aligned via Procrustes. We then correlated parcel-wise gradient values between datasets using Spearman’s r (Fig. 6b). Replicability was strongest for G1 (*r* = 0.74, *p* = 0), and moderate for G2 (*r* = 0.28, *p* = 0.04) and G3 (r = 0.40, p = 0), confirming that G1 is the most stable and reproducible axis, with G2/G3 showing moderate but reliable cross-dataset correspondence.

## Discussion

One of the central goals of systems neuroscience is to derive compact, biologically interpretable coordinates that summarize distributed brain dynamics while respecting anatomical and physiological constraints. Low-dimensional maps—whether derived from connectivity, transcriptomics, or microstructure—have linked mesoscale activity to cortical hierarchy, development, and disease (Bethlehem et al. 2020; Pang et al. 2023; Huntenburg et al. 2018; Sydnor et al. 2021; Margulies et al. 2016a). What has long been missing is an electrophysiology-based representation that preserves temporal specificity and bridges oscillatory mechanisms with macroscale structure. By constructing neurophysiological gradients from source-localized resting-state MEG, we provide precisely such a representation: a stable, low-dimensional embedding of cortical spectral profiles with clear biological anchors and clinical implications.

The first three gradients explained substantial variance in spectral similarity and mapped systematically onto Yeo networks, mirroring the canonical unimodal–transmodal axis seen in fMRI (Margulies et al. 2016b; Paquola et al. 2020). Conceptually, this indicates that, despite the temporal complexity of oscillations, resting-state activity exhibits smooth spatial organization, likely constrained by myeloarchitecture, long-range connectivity, and cytoarchitectonics (Glasser and Essen 2011; Huntenburg et al. 2018). Methodologically, extending gradient analysis to source-space MEG provides a time-resolved complement to hemodynamic gradients and a platform for joint modeling of electrophysiology and fMRI (Shafiei et al. 2023; Vidaurre et al. 2018). High test–retest reliability and cross-dataset replication—most pronounced for G1—suggest that these axes behave as trait-like features suitable for biomarker development (Noble et al. 2021; Hong et al. 2020).

The strong agreement between MEG gradients and geometric/structural MRI gradients anchors oscillatory organization in established macroscale scaffolds. The alignment of G1 with cortical geometry and structural connectivity accords with evidence that folding patterns and myelin content shape large-scale communication and hierarchy (Burt et al. 2018; Huntenburg et al. 2018; Glasser and Essen 2011) . Our results further show that spectral fingerprints (e.g., α/β dominance and aperiodic components) also follow these axes, implying common generative constraints spanning anatomy, conduction delays, and synaptic physiology (Buzsáki and Draguhn 2004; Deco et al. 2013). Practically, this cross-modal convergence supports using neurophysiological gradients as integration coordinates for multimodal connectomics and cross-modal inference.

Embedding a Wilson–Cowan neural-mass model into empirical connectomes shows that variation along cortical gradients is mechanistically consistent with changes in local excitatory–inhibitory (E–I) balance. Specifically, the second principal gradient (G2) is negatively correlated with the simulated E/I ratio, defined here as the time-averaged excitatory-to-inhibitory activity in the model. This pattern aligns with theory and data indicating that E–I balance shapes oscillatory spectra and modulates neuronal response (Brunel and Wang 2003; Gao et al. 2017; Deco et al. 2013). Importantly, we show that these circuit-level parameters map onto macroscale spatial organization: regional spectral profiles covary with both local microcircuit properties and each region’s network embedding in the connectome. This establishes a basis for model-driven phenotyping of conditions hypothesized to involve E–I dysregulation (e.g., schizophrenia spectrum, autism, neurodegeneration). Neurophysiological gradients co-localize with PET-derived neurotransmitter receptor maps (G1 particularly with 5-HT and GABA; G2/G3 with dopaminergic/glutamatergic systems), with BigBrain histological/microstructural gradients, and with cell-type gene-expression maps. These results extend prior links between receptor density and functional networks and between histological gradients and cortical hierarchy (Paquola et al. 2020; Hansen and Misic 2025b; Hansen et al. 2022b), suggesting that oscillatory axes arise from multi-level constraints: receptor landscapes set regional time constants and gain, laminar architecture shapes intrinsic coupling and resonant frequencies, and cell-type composition tunes inhibitory/excitatory balance—together yielding the spectral “fingerprints” captured by the gradients (Schulz et al. 2015; Bastos et al. 2012).

Viewed through the lens of gradients, aging appears as a continuous spatial reweighting, consistent with age effects on α/β power, 1/f slope, and network segregation, while offering a spatially continuous organizational perspective (Voytek et al. 2015; Sala-Llonch et al. 2015). Projecting these changes onto cognitive domains (language, memory, multisensory integration) provides a new way to contextualize lifespan brain dynamics and to develop biomarkers for healthy and pathological aging. In PD, we observe significant gradient alterations—particularly in regions linked to language/memory, social cognition, and multisensory processing—consistent with widespread oscillatory abnormalities and β-band disturbances (Hirano and Uhlhaas 2021; Olde Dubbelink et al. 2014; Hammond et al. 2007).

Framing PD within gradient space yields a spatially continuous, mechanistically interpretable phenotype that bridges local β-pathophysiology with network-level dysfunction. This framework may facilitate subtype stratification, guide network-targeted neuromodulation (e.g., adaptive DBS or stimulation along gradient axes), and provide parsimonious outcome measures for clinical trials.

### Limitation

Several limitations of this study should be noted. First, the work primarily relied on resting-state MEG, structural MRI and functional MRI data, while other modalities such as task-evoked electrophysiology or high-resolution laminar imaging were not included; thus, the generalizability of the gradients to diverse brain states remains unclear. Second, some gradients or frequency bands exhibited modest correlations, potentially due to measurement noise (sensor noise, head-position variability), state fluctuations (arousal, mind wandering), uncertainties in spectral parameterization for weak or overlapping peaks, and preprocessing differences across sessions/datasets. These factors do not undermine the utility of the gradients but constrain the sensitivity for detecting individual differences. Future work could improve reliability by incorporating hierarchical measurement models, accounting for peak-detection uncertainty, and applying individualized head-position corrections. Third, while PD was used as a clinical model, the sample size was relatively modest and limited to mild-to-moderate stages, restricting the ability to generalize findings across disease heterogeneity. Finally, demographic and lifestyle diversity was limited in the datasets, and potential confounds such as medication effects could not be fully ruled out. Future studies should expand to more heterogeneous cohorts, integrate longitudinal and task-based paradigms, and test gradient alterations across multiple disorders to strengthen both mechanistic and translational insights.

## Conclusion

Our findings integrate electrophysiological, structural, molecular, and clinical perspectives into a unified framework of neurophysiological gradients. By demonstrating their stability, biological grounding, age sensitivity, and clinical alterations, we establish gradients as a powerful systems-level tool for bridging basic neuroscience and translational applications. Future research should expand this framework to other disorders, integrate with multimodal datasets, and explore longitudinal trajectories, ultimately paving the way for gradient-based biomarkers of brain health and disease.

## Method

### Exploration dataset

#### Cambridge Centre for Aging and Neuroscience Project Dataset

In this study, we included 608 healthy adult participants (307 males, 301 females; mean age = 54.19 ± 18.19 years) from the Cambridge Centre for Ageing and Neuroscience (Cam-CAN) dataset (Taylor et al. 2017), all of whom underwent both resting-state magnetoencephalography (MEG) and magnetic resonance imaging (MRI).

#### MEG Data Acquisition

Resting-state MEG recordings were conducted at the MRC Cognition and Brain Sciences Unit (MRC-CBSU, Cambridge) using a 306-channel VectorView MEG system (Elekta Neuromag, Helsinki), comprising 102 magnetometers and 204 orthogonal planar gradiometers housed within a magnetically shielded room (MSR). Data were sampled at 1000 Hz and filtered online using a 0.03–330 Hz band-pass filter. Continuous tracking of head position within the MEG helmet was enabled by four Head Position Indicator (HPI) coils, allowing for offline motion correction. Vertical and horizontal electrooculogram (VEOG, HEOG) signals were recorded using bipolar electrodes to monitor eye blinks and saccades, while a separate bipolar electrode pair recorded electrocardiogram (ECG) signals to detect pulse-related artifacts. The MEG protocol included approximately 8 minutes and 40 seconds of resting-state data collection.

#### MRI Data Acquisition

Structural and functional MRI data were acquired during a separate scanning session at the same research facility. High-resolution T1-weighted anatomical images and Diffusion-weighted imaging (DWI) data were collected for cortical surface reconstruction and atlas-based registration. Resting-state functional MRI (rs-fMRI) data were acquired using a gradient-echo echo-planar imaging (EPI) sequence with the following parameters: repetition time (TR) = 1970 ms, echo time (TE) = 30 ms, flip angle = 78°, field of view (FOV) = 192 × 192 mm², voxel size = 3 × 3 × 4.44 mm³, and 32 axial slices covering the cerebellum. Each rs-fMRI run lasted approximately 8 minutes and 40 seconds. An additional multi-echo EPI sequence was acquired for improved signal-to-noise ratio (TR = 2470 ms; TEs = 9.4, 21.2, 33, 45, and 57 ms) using the same spatial resolution and slice coverage. To correct for magnetic susceptibility-induced distortions, dual-echo gradient-echo field maps were acquired using a spoiled gradient recalled echo (SPGR) sequence (TE1 = 5.19 ms, TE2 = 7.65 ms), with a total acquisition time of 54 seconds.

### Replication dataset

#### The Open MEG Archives

This study utilized resting-state MEG data obtained from the Open MEG Archives (OMEGA) (Niso et al. 2016), all acquired using a uniform MEG system (275-channel whole-head CTF; Port Coquitlam, British Columbia, Canada). Recordings were sampled at 2400 Hz with a low-pass anti-aliasing filter set at 600 Hz and employed a built-in third-order spatial gradient noise cancellation protocol.

Resting-state MEG data from 158 unrelated participants (77 females; mean age: 31.9 ± 14.7 years) were analyzed. Each session comprised approximately five minutes of continuous recording. A subset of these individuals (N = 47) underwent multiple recording sessions across separate days and were included in the longitudinal fingerprinting analysis. The data acquisition and management procedures were approved by the Research Ethics Board of the Montreal Neurological Institute, and all research activities adhered to the ethical guidelines of the same institution.

#### MEG Data Preprocessing

MEG data preprocessing and analysis were performed using *Brainstorm* (2016 version) (Tadel et al. 2011) with default parameters unless otherwise specified, following established best-practice guidelines. Initial preprocessing included the removal of slow-wave and DC-offset components via a high-pass finite impulse response (FIR) filter with a cutoff frequency of 0.3 Hz. Raw MEG signals were visually inspected to identify and exclude bad channels and artifact-laden time segments. Power-line noise at 50 Hz and its harmonics were attenuated using notch filtering. A subsequent low-pass filter (0.6–280 Hz) was applied to suppress high-frequency noise while retaining the relevant neural signal. Physiological artifacts, including cardiac and ocular artifacts, were removed using signal-space projection (SSP) techniques, with artifact components identified based on electrocardiogram (ECG) and electrooculogram (EOG) recordings.

Preprocessed MEG data were segmented into 30-second non-overlapping epochs to ensure the robustness of resting-state analyses. Segments were band-pass filtered to the frequency range of interest and resampled at four times the upper cutoff frequency to prevent aliasing while preserving signal fidelity.

Source reconstruction was conducted using each participant’s T1-weighted anatomical MRI, acquired on a 1.5T Siemens Sonata scanner. Cortical surface reconstruction and tissue segmentation were performed using *FreeSurfer* (Fischl 2012). Cortical surfaces were tessellated into triangular meshes and visualized using inflated surface renderings. Co-registration of MEG sensor positions to individual anatomy was accomplished using digitized scalp landmarks obtained during the MEG session. Forward modeling was carried out using the overlapping spheres approach, and cortical source activity was estimated via linearly constrained minimum variance (LCMV) beamforming. Noise covariance matrices, derived from empty-room recordings, were used for normalization to correct for depth bias in source estimation.

#### Neurophysiological Gradient Construction

The whole-cortex power spectrum was computed at the vertex level and parcellated using the Schaefer 200-region atlas. Regional spectral sequences were then extracted using principal component analysis (PCA). To quantify the similarity of neural oscillatory dynamics across cortical regions, we computed the covariance matrix of regional spectral sequences. Similarly, whole-brain neurophysiological gradients were computed using the BrainSpace toolbox (v0.1.10; https://github.com/MICA-MNI/BrainSpace) with default parameter settings (Vos de Wael et al. 2020). The covariance matrix was first z-transformed, and only the top 10% of the strongest connections per region were retained, following established procedures. An affinity matrix based on cosine similarity was then constructed to quantify the similarity in oscillatory profiles across regions. Subsequently, diffusion map embedding, a nonlinear dimensionality reduction technique, was applied to extract low-dimensional manifold representations from the high-dimensional covariance data at the individual level. The first three gradient components were retained for downstream analyses. To ensure cross-subject comparability, individual gradients were aligned to a reference template derived from the group-averaged oscillatory covariance matrix using Procrustes alignment from respective dataset.

#### fMRI Data Preprocessing and Functional Gradient Construction

Functional MRI data were first converted from raw DICOM format to the Brain Imaging Data Structure (BIDS) (Gorgolewski et al. 2016) format using *HeuDiConv* v0.13.1. Subsequent structural and functional preprocessing was performed using *fMRIPrep* version 23.0.2 (Esteban et al. 2019), built upon the *Nipype* 1.8 framework (Gorgolewski et al. 2011). Anatomical preprocessing steps included intensity normalization, skull stripping, tissue segmentation, cortical surface reconstruction, and spatial normalization to MNI space. Functional preprocessing comprised motion correction, slice-timing correction, and co-registration to the corresponding T1-weighted anatomical image. Following preprocessing, the functional time series were parcellated according to the Schaefer 200 × 7 cortical atlas. Confound regression was conducted using the “simple” strategy, which included high-pass filtering, removal of motion-related and tissue-based signals, linear detrending, and z-score normalization. Functional connectivity matrices were then constructed for each participant by computing zero-lag Pearson correlation coefficients between regional mean time series.

Whole-brain functional connectivity gradients were computed using the BrainSpace toolbox with the same parameters as used in MEG gradient. To improve the reliability of gradient estimation, functional connectivity matrices were first z-scored, and only the top 10% of strongest connections per region were retained in line with established preprocessing protocols. A cosine similarity-based affinity matrix was constructed to capture the similarity between regional connectivity profiles. Diffusion map embedding (Coifman and Lafon 2006) was then applied to extract low-dimensional representations of individual-level connectomes. We computed the first ten gradient components and selected the principal gradient, corresponding to the canonical sensory-to-association axis, for subsequent analyses. To facilitate inter-subject comparability, individual gradients were aligned to a group-level template generated from 100 unrelated participants in the Human Connectome Project using Procrustes transformation (Vos de Wael et al. 2020).

#### Diffusion MRI Preprocessing and Structural Gradient Construction

DWI data from CAM-CAN were preprocessed using *QSIprep* version 0.17.0 (Cieslak et al. 2021). The preprocessing pipeline included denoising, Gibbs ringing artifact removal, bias field correction, intensity normalization, head motion and eddy current correction, as well as image registration and normalization to standard space. Structural connectivity reconstruction was conducted using the “mrtrix_multishell_msmt_ACT-hsvs” pipeline. Fiber orientation distributions were estimated using multi-shell multi-tissue constrained spherical deconvolution (MSMT-CSD) (Jeurissen et al. 2014). Probabilistic tractography was performed using the iFOD2 algorithm with anatomically constrained tractography (ACT), incorporating T1-weighted segmentation priors. Streamline weights were computed using the SIFT2 algorithm, and weighted structural connectivity matrices were generated accordingly(Smith et al. 2015).

Similarly, whole-brain structural connectivity gradients were computed using the BrainSpace toolbox with default parameter settings. The structural connectivity matrix was first z-transformed, and only the top 10% of the strongest connections per region were retained, in accordance with established procedures in the literature. An affinity matrix was then constructed based on cosine similarity to quantify the similarity of connectivity profiles across regions. Subsequently, diffusion map embedding, a nonlinear dimensionality reduction technique, was applied to derive low-dimensional manifold representations from high-dimensional structural connectome data at the individual level. The first three gradient components were extracted for downstream analyses. To ensure cross-subject comparability, individual gradients were aligned to a reference template constructed from the group-averaged structural connectivity matrix across all participants using Procrustes alignment.

#### Structural MRI Preprocessing and Geometric Gradient Construction

High-resolution T1-weighted anatomical MRI scans were processed using *FreeSurfer* with the standard *recon-all* pipeline, including skull stripping, intensity normalization, surface reconstruction, and cortical parcellation. Individual white matter surfaces were extracted as triangulated meshes, resampled to ∼164,000 vertices, and aligned to the *fsaverage* template via spherical registration. These standardized surface meshes were used as the geometric domain for Laplace–Beltrami eigenmode computation (Pang et al. 2023). Quality control was performed to ensure accurate surface topology.

To characterize the intrinsic geometric structure of the cortical surface, we computed geometric eigenmodes by solving the Laplace–Beltrami operator (LBO) on the cortical manifold (Pang et al. 2023). The LBO was discretized over the standard *fsaverage* surface mesh from *FreeSurfer*, containing approximately 164,000 vertices. The LBO matrix was constructed using a cotangent-weighted scheme applied to the mesh’s triangular faces. Eigenmodes were obtained by solving the Helmholtz equation Δψ = −λψ with Neumann boundary conditions, where Δ denotes the discrete LBO, λ the eigenvalues, and ψ the corresponding eigenfunctions. These eigenfunctions represent a hierarchy of spatially smooth basis functions defined by the geometry of the cortical surface, ordered from low to high spatial frequency. The first non-constant eigenmode corresponds to the smoothest spatial gradient, while higher-order modes encode increasingly fine-grained spatial variations. In this study, we retained the first *N* = 200 non-trivial eigenmodes to provide a multiscale functional basis for projecting and analyzing cortical maps. This geometric framework enables biologically interpretable decompositions of functional or structural data and has the advantage of being anatomically grounded, spatially orthogonal, and inherently multiscale.

#### Wilson Cowan Modeling

The Wilson–Cowan (WC) model is a classical framework for describing large-scale brain network dynamics (Wilson and Cowan 1972). It consists of two interacting neuronal populations—excitatory and inhibitory—that are linked through coupling parameters(Wilson and Cowan 1972). The dynamics of each population include several components: local coupling terms reflecting excitatory and inhibitory interactions, a time-delayed network interaction term determined by the structural connectivity and transmission delays, a threshold and nonlinear response implemented via a sigmoid transformation, and modulation by refractory effects together with a time constant governing the rate of change.

In this study, we first extracted empirical brain activity features from MEG recordings. Specifically, we parameterized neural power spectra to estimate the center frequency of each brain region (Donoghue et al. 2020). Center frequency was selected as the fitting feature because it is interpretable and relatively robust to noise, being sensitive to local excitatory–inhibitory dynamics while also reflecting network influences. The WC model was embedded in individualized structural connectivity (Sanz Leon et al. 2013). The estimated center frequency therefore served as the objective feature for fitting the model. Individual structural connectivity data provided the network topology for embedding the WC model into the brain connectome (Sanz Leon et al. 2013). By tuning parameters such as global coupling strength and conduction velocity, the model generated region-wise predictions of center frequency (Bansal et al. 2019). Model accuracy was quantified as the correlation between simulated and empirical vectors of regional center frequencies; accordingly, the optimization targeted the spatial distribution of center frequencies rather than their absolute values, which would be emphasized by objectives such as *Pearson’s* correlation. Details of the fitting procedure are provided in the *Supplementary Material* (“*Parameter Setting for Wilson–Cowan Modeling*”).

#### Genetic analysis and enrichment analysis

To investigate transcriptional mechanisms underlying large-scale network organization, we estimated cell-type proportions by deconvolving microarray expression data from the Allen Human Brain Atlas (AHBA; http://human.brain-map.org/) (Hawrylycz et al. 2012). The AHBA contains gene expression profiles from six neurotypical adult postmortem brains (mean age = 42.5 ± 13.4 years; 5 males, 1 female), comprising 3,702 spatially distinct cortical tissue samples. To spatially align transcriptomic data with neuroimaging-based parcellations, we used the abagen toolbox ((Markello et al. 2021); https://github.com/netneurolab/abagen) to preprocess the microarray data and map it to the Schaefer 400 atlas. We then examined correlations between neural timescale statistical maps and transcriptionally dysregulated messenger RNA (mRNA) expression profiles, focusing on genes with elevated expression in cortical regions. To identify convergent and divergent biological pathways involved in functional reorganization, we performed gene enrichment analysis using Metascape (https://metascape.org; (Zhou et al. 2019)), an integrative resource drawing from over 40 biological databases. Genes most strongly associated with patterns of functional reorganization were submitted for enrichment analysis. Statistical significance was defined as false discovery rate (FDR)-corrected q < 0.05, and results were validated against a null distribution.

#### Cortical annotations

Furthermore, to physiologically annotate the cortical organization of the brain’s magnetic gradients, we linked these gradients with normative maps of BigBrain, cell types, and neurotransmitter receptors. BigBrain intensity profiles were generated using the BigBrainWarp toolbox (Amunts et al. 2013; Paquola et al. 2021) to quantify cytoarchitectural features based on cell staining intensity, calculated across 50 equivolumetric surfaces (Wagstyl et al. 2020; Hawrylycz et al. 2012; Wagstyl et al. 2018).

In this study, the brain’s magnetic gradients were correlated with 24 cellular maps reported in a previous study by (Zhang et al. 2025). Molecular signature profiles for all cell classes were constructed from snDrop-seq samples provided by (Jorstad et al. 2023). Subsequently, cell type proportions were estimated via deconvolution of microarray samples obtained from the AHBA (Hawrylycz et al. 2012). The 24 cell types included Lamp5, Pax6, Vip, Sncg, Lamp5, Lhx6, L5ET, L5/L6 NP, L6 CT, L6b, Astro, VLMC, Endo, Micro/PVM, Oligo, OPC, L2/3 IT, L6 IT Car3, L4 IT, L6 IT, L5 IT, Chandelier, Pvalb, Sst, and Sst Chodl.

Neurotransmitter receptor and transporter densities were quantified using positron emission tomography (PET) tracer maps encompassing 18 molecular targets across nine major neurotransmitter systems, curated by (Hansen et al. 2022b)(https://github.com/netneurolab/hansen_receptors). These systems included dopamine (D₁, D₂, DAT), norepinephrine (NET), serotonin (5-HT₁A, 5-HT₁B, 5-HT₂, 5-HT₄, 5-HT₆, 5-HTT), acetylcholine (α4β2, M₁, VAChT), glutamate (mGluR₅), γ-aminobutyric acid (GABAₐ), histamine (H₃), cannabinoid (CB₁), and opioid (MOR), consistent with prior neurochemical mapping work (Markello and Misic 2021). PET images were nonlinearly registered to the MNI-ICBM 152 (2009c asymmetric) standard space and subsequently parcellated according to the Schaefer 200 cortical atlas. For receptors or transporters with multiple PET maps derived from the same tracer (e.g., 5-HT₁B, D₂, VAChT), weighted averaging was applied to generate a single representative map per target.

#### Null model

In this study, we aimed to quantify the topographic correspondence between neural timescale abnormalities and other neurobiological features. To determine the statistical significance of these spatial associations while controlling for inherent spatial autocorrelation, we implemented a spatial permutation–based null model. First, receptor maps were spatially aligned with the neural timescale statistical map, and their correlations were computed. To generate the null distribution, we applied spherical rotations to the cortical surface, which preserved local spatial structure but disrupted the original spatial alignment. Node values were reassigned according to the nearest rotated parcel, and this process was repeated 1,000 times (Markello and Misic 2021). Rotations were performed on one hemisphere and mirrored to the opposite hemisphere to preserve interhemispheric symmetry. The observed correlation was then compared to the null distribution, with statistical significance defined as exceeding the 95*th* percentile of null correlations obtained from both spatial and temporal permutation models.

#### Age effect

Furthermore, we leveraged the Cam-CAN dataset to investigate lifespan-wide reorganization patterns of physiological gradients. Specifically, we performed Pearson‘s correlation with FDR correction analyses between age and each of the three derived physiological gradients, with handedness and sex included as covariates. Also, we further assessed the cognitive relevance of age effect patterns using Neurosynth (Yarkoni et al. 2011). Specifically, we drew on twenty cognition-related meta-analytic maps from prior work (Margulies et al. 2016)—spanning sensory processes to higher-order cognition—and computed Pearson correlations between these maps and the age effect on neurophysiological-gradient maps .

#### Test–retest reliability assessment analysis

To evaluate the test–retest reliability of the neurophysiological gradient, we calculated the intraclass correlation coefficient (ICC) according to its definition. From this definition, it is evident that test–retest reliability integrates both within-subject and between-subject variability. Low within-subject variability or high between-subject variability leads to higher test–retest reliability. Since reliability constrains the validity of various indices used in clinical diagnosis, it represents a fundamental requirement for the development of biomarkers in applied settings. Here, we computed the correlation between the neurophysiological gradient derived from the OMEGA dataset and those obtained from the physiological gradient and CAM-CAN.

## DATA AVAILABILITY

MEG, fMRI, DWI data could be available on https://cam-can.mrc-cbu.cam.ac.uk/dataset/. OMEGA dataset could be available on https://www.mcgill.ca/bic/neuroinformatics/omega. Public resources used in this study can be accessed online, including the Allen Human Brain Atlas (AHBA; http://human.brain-map.org/), neuromaps (https://netneurolab.github.io/neuromaps/usage.html), BrainSpace (https://brainspace.readthedocs.io/en/latest/), and the ENIGMA Toolbox (https://enigma-toolbox.readthedocs.io/en/latest/pages.html). The Characterization and Treatment of Adolescent Depression (CAT-D) dataset is available via OpenNeuro (https://openneuro.org/datasets/).

## CODE AVAILABILITY

Code will be available upon reasonable request.

## ACKNOWLEDGMENTS

Xiaobo Liu is supported by the China Scholarship Council.

## The parameter setting for Wilson Cowan Modeling

In the structural connectivity and preprocessing stage, we used the Schaefer-200 parcellation–based structural connectome. The fiber-count matrix was max-normalized and served as the interregional weight matrix, whereas the mean fiber-length matrix quantified interregional distances. The primary analyses were conducted on individual connectomes.

For the model and interregional interactions, we adopted a Wilson–Cowan two-population node model to describe local dynamics within each brain region. Each node explicitly included two types of local coupling terms: one capturing within-population (self–self) interactions and the other capturing cross-population (self–other) interactions. Long-range inputs were implemented as time-delayed network interactions: inputs from other regions were weighted by the structural connectivity and propagated with transmission delays determined by fiber lengths and conduction velocity. Both excitatory and inhibitory populations received these long-range inputs.

Regarding parameters, search space, and external drive, the excitatory local gain was treated as a free parameter in the range 0.8–1.2; the inhibitory gain was fixed to one quarter of the excitatory gain; the delay/time-constant parameter ranged from 12 to 14; and the global coupling (network gain) ranged from 12 to 14. External drives to the two populations were set to the four-tuple [1, 0.75, 1, T], where 0.75 is an amplitude scaling factor and T is the total simulation duration; this drive was applied identically to all 200 nodes. Parameter fitting used a uniformly sampled random grid.

For numerical simulations, each parameter set was run on the whole-brain network to obtain excitatory time series for all nodes. The integration step was 0.5 ms for a total of 30,000 steps; the sampling rate followed the implementation and no additional resampling was performed. Power spectral density was computed for each node’s excitatory time series, and the peak frequency within 4–30 Hz was taken as the node’s center frequency to form the simulated center-frequency vector. The empirical center-frequency vector was derived from MEG data.

For the objective function and fitting strategy, we targeted the spatial distribution of center frequencies, using the Pearson correlation between empirical and simulated center-frequency vectors as the fitting score. This emphasizes spatial concordance and robustness rather than minimizing per-node amplitude errors. We performed a random grid search with 500 parameter samples; after simulation and feature extraction for each sample, we computed the correlation and selected the parameter set with the highest score as the optimum, recording both the optimal parameters and the corresponding fit.

**Supplementary Fig. 1.**
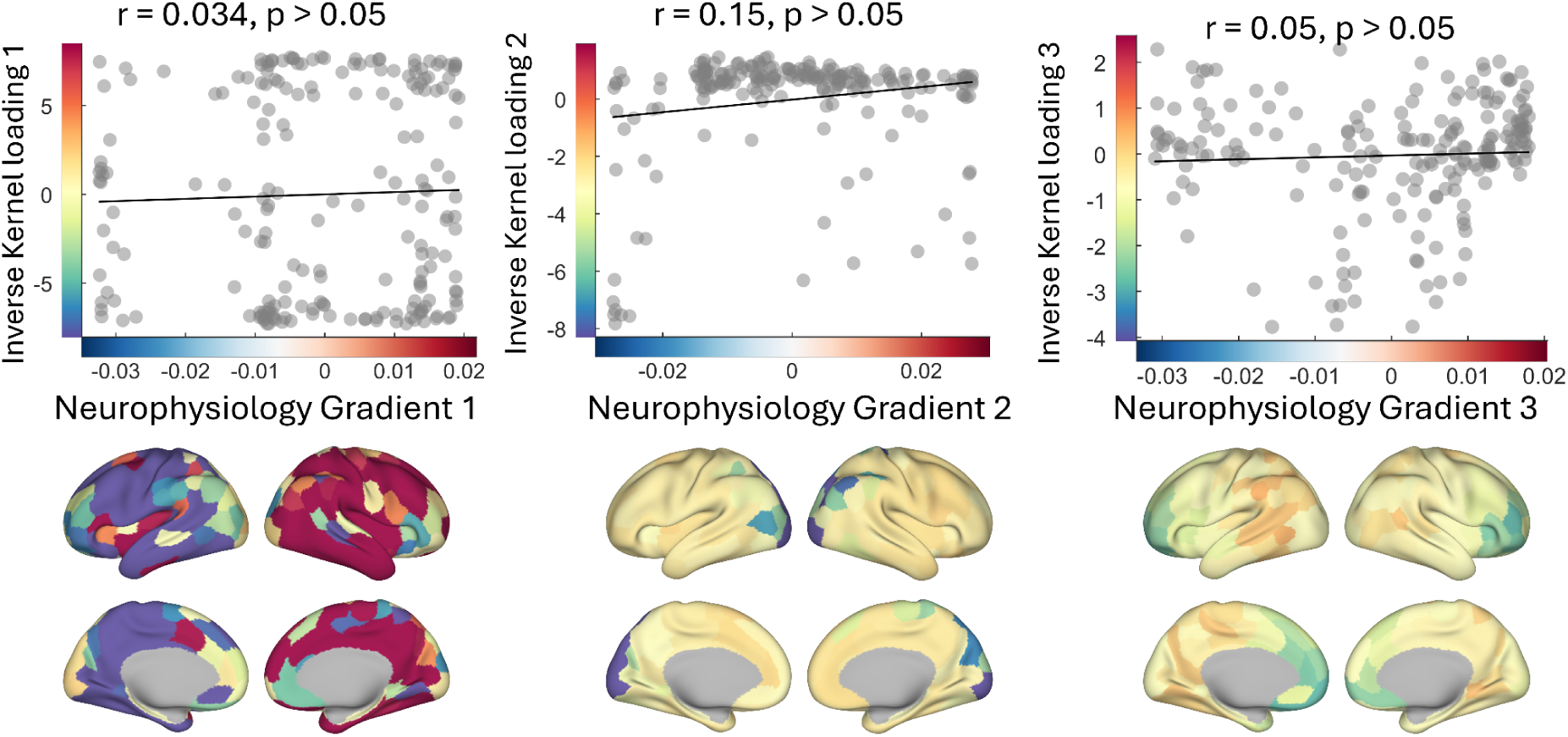
Relationships between neurophysiological gradients and inverse kernel loadings, and their spatial distributions. The top row presents scatterplots illustrating the linear associations between Neurophysiology Gradients 1–3 and the corresponding inverse kernel loadings (y-axis); black lines indicate ordinary least-squares fits. All correlations are weak and not statistically significant: Gradient 1 (*r* = 0.034, *p* > 0.05), Gradient 2 (*r* = 0.15, *p* > 0.05), and Gradient 3 (*r* = 0.05, *p* > 0.05).

**Supplementary Fig. 2.**
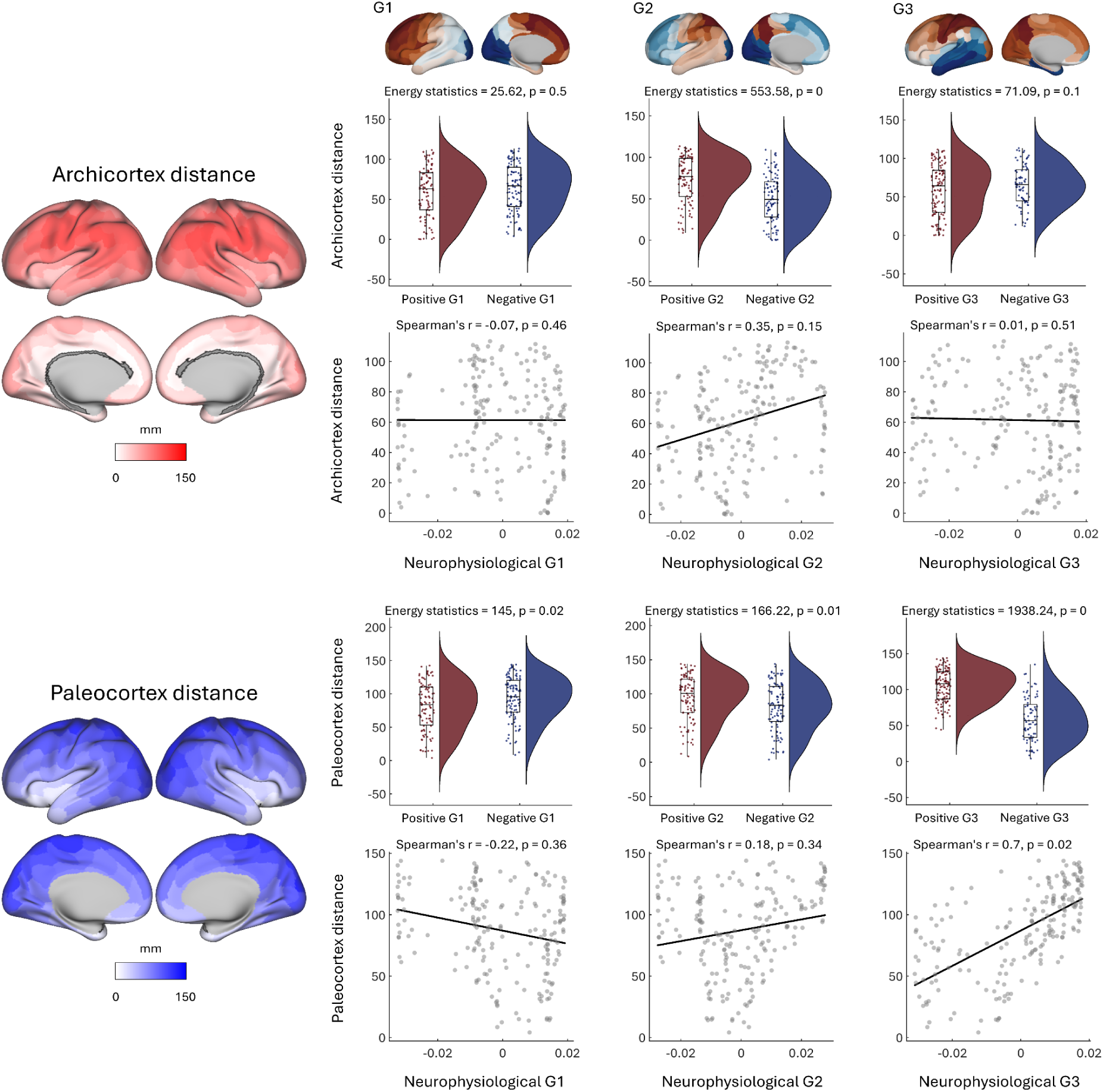
Relationship between MEG neurophysiological gradients and distance to the archi-/paleocortex under the dual-origin theory. Left: geodesic distances to two evolutionarily ancient cortical origins rendered on a standard cortical surface—top, distance to the archicortex; bottom, distance to the paleocortex (color bar in mm). Right: three columns correspond to the three neurophysiological gradients (G1–G3). In each column, the top row shows violin/box plots comparing distance distributions between regions with positive versus negative gradient values, reporting the energy statistic and its significance; the bottom row shows scatterplots with Spearman correlations (x-axis: the corresponding neurophysiological gradient; y-axis: distance to the archi-/paleocortex). Results: paleocortical distance increases with gradient strength and is significantly associated with G2–G3, with clear separations between positive and negative gradient regions; archicortical distance also shows a significant separation for G2, whereas effects for G1 and G3 are weaker. Overall, the findings support the dual-origin theory’s prediction that cytoarchitectonic and topological constraints originating from the paleo/archicortex are reflected in magnetoencephalographic neurophysiological gradients.

**Supplementary Fig. 3.**
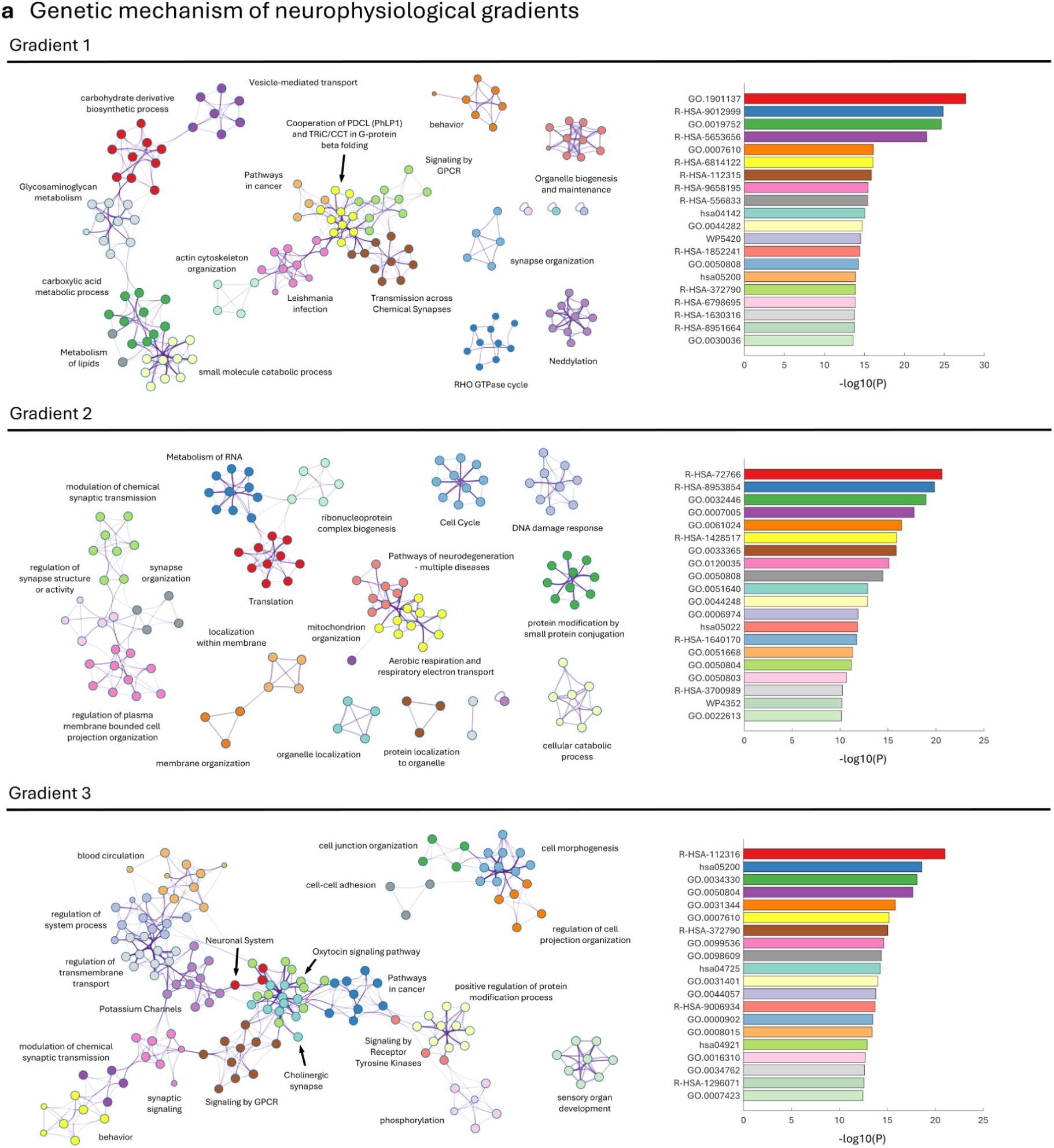
Genetic enrichment analysis of neurophysiological gradients. Panels are arranged by gradient (Gradients 1–3). For each gradient, the left panel shows a gene-set enrichment network: nodes denote significantly enriched pathways/GO terms (sourced from Reactome/GO; node colors indicate modules/clusters, node size approximates enrichment strength or gene count), and edges represent gene overlap or semantic similarity between terms; arrows mark representative hub pathways. The right panel lists the top enriched pathways ranked by −log10(P). Gradient 1 is enriched for synaptic transmission and signaling (e.g., GPCR signaling), vesicle-mediated transport, cytoskeletal/cell-organelle biogenesis and maintenance, and metabolic processes (e.g., lipid and carbohydrate metabolism). Gradient 2 highlights protein synthesis and ribosomal/ribonucleoprotein assembly, mitochondrial organization and aerobic respiration/electron transport, RNA metabolism and translational regulation, the cell cycle and DNA damage response, as well as synapse organization and function. Gradient 3 is enriched for nervous-system–related processes (ion/potassium channels, chemical synaptic transmission, GPCR and oxytocin signaling), cell adhesion and junction organization, receptor tyrosine kinase signaling, and pathways underlying morphogenesis and projection organization.

**Supplementary Fig. 4.**
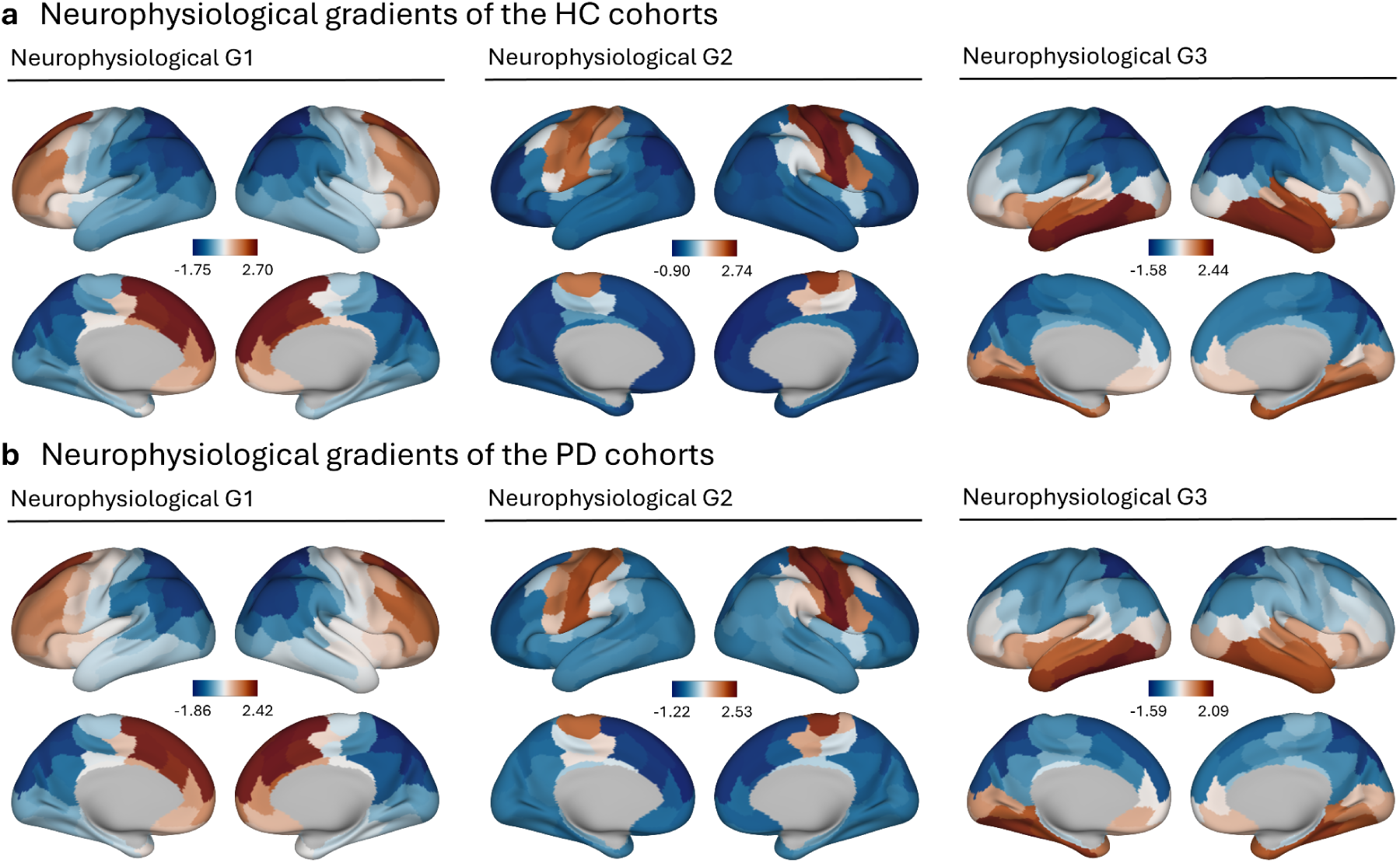
Neurophysiological gradients (G1–G3) in the HC and PD cohorts. (a) Healthy control (HC) cohort; (b) Parkinson’s disease (PD) cohort. Each column shows cortical projections of neurophysiological gradients G1, G2, and G3, with lateral and medial views of both hemispheres. The color bar indicates gradient values from low (blue) to high (orange); gray denotes masked/absent data. Compared with HC, the PD cohort exhibits altered gradient distributions across multiple cortical regions, suggesting a reorganization of large-scale functional architecture.

### Parkinson’s disease dataset

## The Open MEG Archives

This study utilized resting-state MEG data with Parkinson’s disease (PD) obtained from the OMEGA, all acquired using a uniform MEG system the same as that used in the replication dataset.

Both healthy controls (n = 50, mean age = 60.58 ± 6.83 years) and patients with idiopathic PD (n = 65, mean age = 64.68 ± 7.93 years) at mild to moderate stages (Hoehn and Yahr scale: 1–3) were recruited through the Quebec Parkinson Network. Participants underwent comprehensive clinical, neuroimaging, neuropsychological, and biological profiling. All PD patients were maintained on a stable regimen of antiparkinsonian medication with satisfactory clinical response prior to study enrollment.

We further computed physiological gradients for the PD and HC groups and employed two-sample t tests to compare physiological gradient reconfiguration between patients and controls.

**Supplementary Fig. 5.**
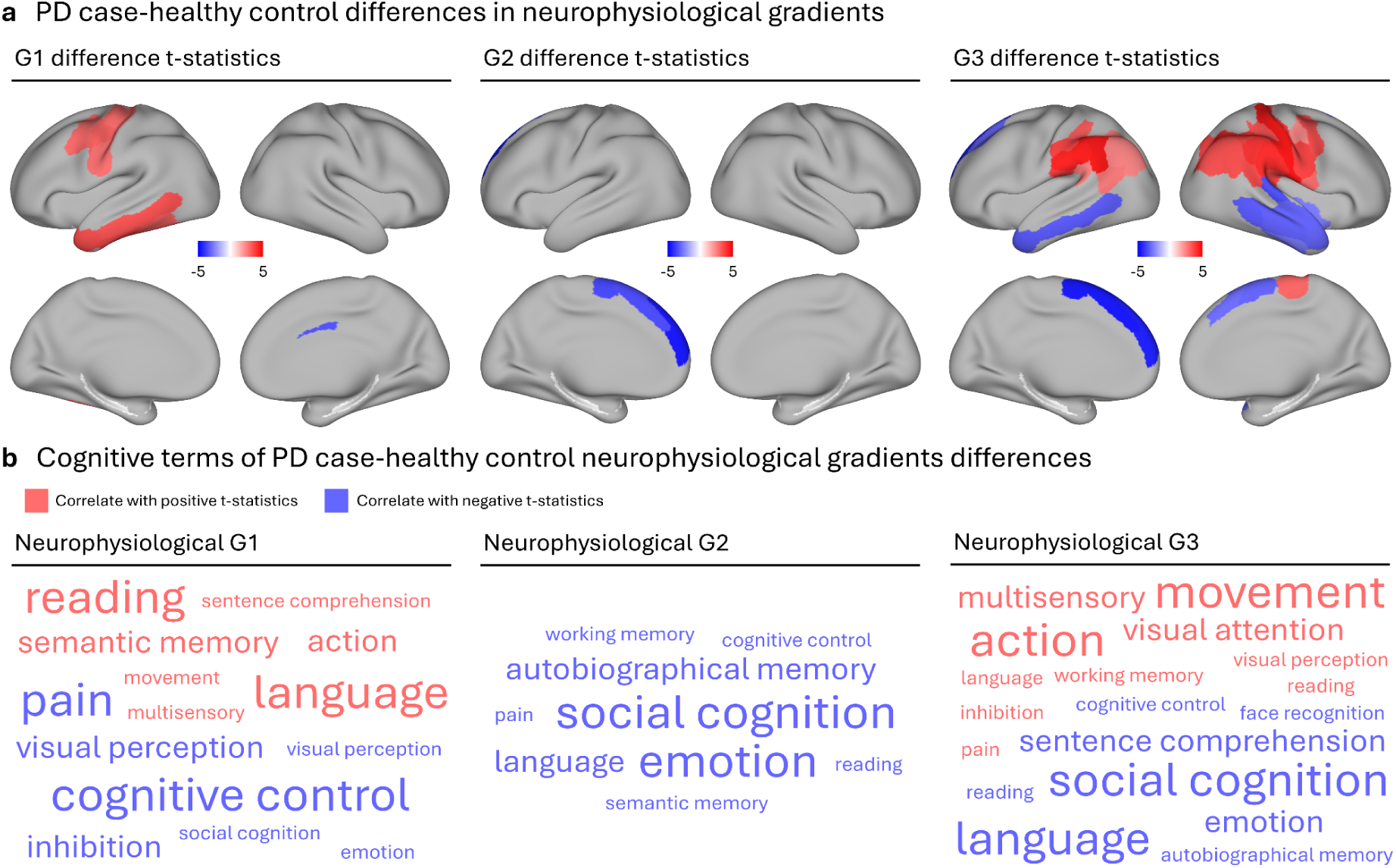
Neurophysiological Gradients Alternation in Parkinson’s Disease. **a.** Cortical projections showing associations between each gradient (G1–G3) and age. Red indicates a positive age effect (increase with age), and blue indicates a negative age effect (decrease with age). **b.** Word clouds representing cognitive functions whose spatial maps correlate with the age-related effects of each gradient. Red terms indicate positive age associations; blue terms indicate negative age associations.

## Reference

Amunts, Katrin, Claude Lepage, Louis Borgeat, et al. 2013. “BigBrain: An Ultrahigh-Resolution 3D Human Brain Model.” Science 340 (6139): 1472–75. 10.1126/science.1235381.

Bansal, Kanika, Javier O. Garcia, Steven H. Tompson, Timothy Verstynen, Jean M. Vettel, and Sarah F. Muldoon. 2019. “Cognitive Chimera States in Human Brain Networks.” Science Advances 5 (4): eaau8535. 10.1126/sciadv.aau8535.

Bastos, Andre M., W. Martin Usrey, Rick A. Adams, George R. Mangun, Pascal Fries, and Karl J. Friston. 2012. “Canonical Microcircuits for Predictive Coding.” Neuron 76 (4): 695–711. 10.1016/j.neuron.2012.10.038.

Belliveau, J. W., D. N. Kennedy, R. C. McKinstry, et al. 1991. “Functional Mapping of the Human Visual Cortex by Magnetic Resonance Imaging.” Science 254 (5032): 716–19. 10.1126/science.1948051.

Beste, Christian, Alexander Münchau, and Christian Frings. 2023. “Towards a Systematization of Brain Oscillatory Activity in Actions.” Communications Biology 6 (1): 137. 10.1038/s42003-023-04531-9.

Bethlehem, Richard A. I., Casey Paquola, Jakob Seidlitz, et al. 2020. “Dispersion of Functional Gradients across the Adult Lifespan.” NeuroImage 222 (November): 117299. 10.1016/j.neuroimage.2020.117299.

Brunel, Nicolas, and Xiao-Jing Wang. 2003. “What Determines the Frequency of Fast Network Oscillations With Irregular Neural Discharges? I. Synaptic Dynamics and Excitation-Inhibition Balance.” Journal of Neurophysiology 90 (1): 415–30. 10.1152/jn.01095.2002.

Burt, Joshua B., Murat Demirtaş, William J. Eckner, et al. 2018. “Hierarchy of Transcriptomic Specialization across Human Cortex Captured by Structural Neuroimaging Topography.” Nature Neuroscience 21 (9): 1251–59. 10.1038/s41593-018-0195-0.

Buzsáki, György, and Andreas Draguhn. 2004. “Neuronal Oscillations in Cortical Networks.” Science 304 (5679): 1926–29. 10.1126/science.1099745.

Cieslak, Matthew, Philip A. Cook, Xiaosong He, et al. 2021. “QSIPrep: An Integrative Platform for Preprocessing and Reconstructing Diffusion MRI Data.” Nature Methods 18 (7): 775–78. 10.1038/s41592-021-01185-5.

Coifman, Ronald R., and Stéphane Lafon. 2006. “Diffusion Maps.” Applied and Computational Harmonic Analysis, Special Issue: Diffusion Maps and Wavelets, vol. 21 (1): 5–30. 10.1016/j.acha.2006.04.006.

Deco, Gustavo, Adrián Ponce-Alvarez, Dante Mantini, Gian Luca Romani, Patric Hagmann, and Maurizio Corbetta. 2013. “Resting-State Functional Connectivity Emerges from Structurally and Dynamically Shaped Slow Linear Fluctuations.” Articles. Journal of Neuroscience 33 (27): 11239–52. 10.1523/JNEUROSCI.1091-13.2013.

Dong, Hao-Ming, Xi-Han Zhang, Loïc Labache, et al. 2024. “Ventral Attention Network Connectivity Is Linked to Cortical Maturation and Cognitive Ability in Childhood.” Nature Neuroscience 27 (10): 2009–20. 10.1038/s41593-024-01736-x.

Donoghue, Thomas, Matar Haller, Erik J. Peterson, et al. 2020. “Parameterizing Neural Power Spectra into Periodic and Aperiodic Components.” Nature Neuroscience 23 (12): 1655–65. 10.1038/s41593-020-00744-x.

Es, Mats W. J. van, Cameron Higgins, Chetan Gohil, Andrew J. Quinn, Diego Vidaurre, and Mark W. Woolrich. 2025. “Large-Scale Cortical Functional Networks Are Organized in Structured Cycles.” Nature Neuroscience 28 (10): 2118–28. 10.1038/s41593-025-02052-8.

Esteban, Oscar, Christopher J. Markiewicz, Ross W. Blair, et al. 2019. “fMRIPrep: A Robust Preprocessing Pipeline for Functional MRI.” Nature Methods 16 (1): 111–16. 10.1038/s41592-018-0235-4.

Fischl, Bruce. 2012. “FreeSurfer.” NeuroImage, 20 YEARS OF fMRI, vol. 62 (2): 774–81. 10.1016/j.neuroimage.2012.01.021.

Gao, Richard, Erik J. Peterson, and Bradley Voytek. 2017. “Inferring Synaptic Excitation/Inhibition Balance from Field Potentials.” NeuroImage 158 (September): 70–78. 10.1016/j.neuroimage.2017.06.078.

Glasser, Matthew F., and David C. Van Essen. 2011. “Mapping Human Cortical Areas In Vivo Based on Myelin Content as Revealed by T1- and T2-Weighted MRI.” Articles. Journal of Neuroscience 31 (32): 11597–616. 10.1523/JNEUROSCI.2180-11.2011.

Godfrey, Megan, and Krish D. Singh. 2021. “Measuring Robust Functional Connectivity from Resting-State MEG Using Amplitude and Entropy Correlation across Frequency Bands and Temporal Scales.” NeuroImage 226 (February): 117551. 10.1016/j.neuroimage.2020.117551.

Gorgolewski, Krzysztof, Christopher D. Burns, Cindee Madison, et al. 2011. “Nipype: A Flexible, Lightweight and Extensible Neuroimaging Data Processing Framework in Python.” Frontiers in Neuroinformatics 5 (August). 10.3389/fninf.2011.00013.

Gorgolewski, Krzysztof J., Tibor Auer, Vince D. Calhoun, et al. 2016. “The Brain Imaging Data Structure, a Format for Organizing and Describing Outputs of Neuroimaging Experiments.” Scientific Data 3 (1): 160044. 10.1038/sdata.2016.44.

Hammond, Constance, Hagai Bergman, and Peter Brown. 2007. “Pathological Synchronization in Parkinson’s Disease: Networks, Models and Treatments.” Trends in Neurosciences 30 (7): 357–64. 10.1016/j.tins.2007.05.004.

Hansen, Justine Y., Ross D. Markello, Jacob W. Vogel, Jakob Seidlitz, Danilo Bzdok, and Bratislav Misic. 2021. “Mapping Gene Transcription and Neurocognition across Human Neocortex.” Nature Human Behaviour 5 (9): 1240–50. 10.1038/s41562-021-01082-z.

Hansen, Justine Y., and Bratislav Misic. 2025a. “Integrating and Interpreting Brain Maps.” Trends in Neurosciences 48 (8): 594–607. 10.1016/j.tins.2025.06.003.

Hansen, Justine Y., and Bratislav Misic. 2025b. “Integrating and Interpreting Brain Maps.” Trends in Neurosciences 48 (8): 594–607. 10.1016/j.tins.2025.06.003.

Hansen, Justine Y., Golia Shafiei, Ross D. Markello, et al. 2022a. “Mapping Neurotransmitter Systems to the Structural and Functional Organization of the Human Neocortex.” Nature Neuroscience 25 (11): 1569–81. 10.1038/s41593-022-01186-3.

Hansen, Justine Y., Golia Shafiei, Ross D. Markello, et al. 2022b. “Mapping Neurotransmitter Systems to the Structural and Functional Organization of the Human Neocortex.” Nature Neuroscience 25 (11): 1569–81. 10.1038/s41593-022-01186-3.

Hawrylycz, Michael J., Ed S. Lein, Angela L. Guillozet-Bongaarts, et al. 2012. “An Anatomically Comprehensive Atlas of the Adult Human Brain Transcriptome.” Nature 489 (7416): 391–99. 10.1038/nature11405.

Hirano, Yoji, and Peter J. Uhlhaas. 2021. “Current Findings and Perspectives on Aberrant Neural Oscillations in Schizophrenia.” Psychiatry and Clinical Neurosciences 75 (12): 358–68. 10.1111/pcn.13300.

Hong, Seok-Jun, Reinder Vos de Wael, Richard A. I. Bethlehem, et al. 2019. “Atypical Functional Connectome Hierarchy in Autism.” Nature Communications 10 (1): 1022. 10.1038/s41467-019-08944-1.

Hong, Seok-Jun, Ting Xu, Aki Nikolaidis, et al. 2020. “Toward a Connectivity Gradient-Based Framework for Reproducible Biomarker Discovery.” NeuroImage 223 (December): 117322. 10.1016/j.neuroimage.2020.117322.

Huntenburg, Julia M., Pierre-Louis Bazin, and Daniel S. Margulies. 2018. “Large-Scale Gradients in Human Cortical Organization.” Trends in Cognitive Sciences 22 (1): 21–31. 10.1016/j.tics.2017.11.002.

Jeurissen, Ben, Jacques-Donald Tournier, Thijs Dhollander, Alan Connelly, and Jan Sijbers. 2014. “Multi-Tissue Constrained Spherical Deconvolution for Improved Analysis of Multi-Shell Diffusion MRI Data.” NeuroImage 103 (December): 411–26. 10.1016/j.neuroimage.2014.07.061.

Jorstad, Nikolas L., Jennie Close, Nelson Johansen, et al. 2023. “Transcriptomic Cytoarchitecture Reveals Principles of Human Neocortex Organization.” Science 382 (6667): eadf6812. 10.1126/science.adf6812.

Kim, Sunghun, Seulki Yoo, Ke Xie, et al. 2024. “Comparison of Different Group-Level Templates in Gradient-Based Multimodal Connectivity Analysis.” Network Neuroscience 8 (4): 1009–31. 10.1162/netn_a_00382.

Kong, Xiaolu, Ru Kong, Csaba Orban, et al. 2021. “Sensory-Motor Cortices Shape Functional Connectivity Dynamics in the Human Brain.” Nature Communications 12 (1): 6373. 10.1038/s41467-021-26704-y.

Kwong, K K, J W Belliveau, D A Chesler, et al. 1992. “Dynamic Magnetic Resonance Imaging of Human Brain Activity during Primary Sensory Stimulation.” Proceedings of the National Academy of Sciences 89 (12): 5675–79. 10.1073/pnas.89.12.5675.

Li, Xiaolu, Shuting Bu, Huize Pang, et al. 2025. “Mapping Striatal Functional Gradients and Associated Gene Expression in Parkinson’s Disease with Continuous Cognitive Impairment.” Npj Parkinson’s Disease 11 (1): 138. 10.1038/s41531-025-01002-2.

Margulies, Daniel S., Satrajit S. Ghosh, Alexandros Goulas, et al. 2016a. “Situating the Default-Mode Network along a Principal Gradient of Macroscale Cortical Organization.” Proceedings of the National Academy of Sciences 113 (44): 12574–79. 10.1073/pnas.1608282113.

Margulies, Daniel S., Satrajit S. Ghosh, Alexandros Goulas, et al. 2016b. “Situating the Default-Mode Network along a Principal Gradient of Macroscale Cortical Organization.” Proceedings of the National Academy of Sciences 113 (44): 12574–79. 10.1073/pnas.1608282113.

Markello, Ross D, Aurina Arnatkeviciute, Jean-Baptiste Poline, Ben D Fulcher, Alex Fornito, and Bratislav Misic. 2021. “Standardizing Workflows in Imaging Transcriptomics with the Abagen Toolbox.” eLife 10 (November): e72129. 10.7554/eLife.72129.

Markello, Ross D., and Bratislav Misic. 2021. “Comparing Spatial Null Models for Brain Maps.” NeuroImage 236 (August): 118052. 10.1016/j.neuroimage.2021.118052.

Meng, Lu, and Jing Xiang. 2016. “Frequency Specific Patterns of Resting-State Networks Development from Childhood to Adolescence: A Magnetoencephalography Study.” Brain and Development 38 (10): 893–902. 10.1016/j.braindev.2016.05.004.

Mostame, Parham, and Sepideh Sadaghiani. 2020. “Phase- and Amplitude-Coupling Are Tied by an Intrinsic Spatial Organization but Show Divergent Stimulus-Related Changes.” NeuroImage 219 (October): 117051. 10.1016/j.neuroimage.2020.117051.

Murray, John D., Alberto Bernacchia, David J. Freedman, et al. 2014. “A Hierarchy of Intrinsic Timescales across Primate Cortex.” Nature Neuroscience 17 (12): 1661–63. 10.1038/nn.3862.

Niso, Guiomar, Christine Rogers, Jeremy T. Moreau, et al. 2016. “OMEGA: The Open MEG Archive.” NeuroImage, Sharing the wealth: Brain Imaging Repositories in 2015, vol. 124 (January): 1182–87. 10.1016/j.neuroimage.2015.04.028.

Noble, Stephanie, Dustin Scheinost, and Robert Todd Constable. 2021. “A Guide to the Measurement and Interpretation of fMRI Test-Retest Reliability.” Current Opinion in Behavioral Sciences, Deep Imaging -Personalized Neuroscience, vol. 40 (August): 27–32. 10.1016/j.cobeha.2020.12.012.

Olde Dubbelink, Kim T. E., Arjan Hillebrand, Diederick Stoffers, et al. 2014. “Disrupted Brain Network Topology in Parkinson’s Disease: A Longitudinal Magnetoencephalography Study.” Brain 137 (1): 197–207. 10.1093/brain/awt316.

Pang, James C., Kevin M. Aquino, Marianne Oldehinkel, et al. 2023. “Geometric Constraints on Human Brain Function.” Nature 618 (7965): 566–74. 10.1038/s41586-023-06098-1.

Paquola, Casey, Jessica Royer, Lindsay B Lewis, et al. 2021. “The BigBrainWarp Toolbox for Integration of BigBrain 3D Histology with Multimodal Neuroimaging.” eLife 10 (August): e70119. 10.7554/eLife.70119.

Paquola, Casey, Jakob Seidlitz, Oualid Benkarim, et al. 2020. “A Multi-Scale Cortical Wiring Space Links Cellular Architecture and Functional Dynamics in the Human Brain.” PLOS Biology 18 (11): e3000979. 10.1371/journal.pbio.3000979.

Paquola, Casey, Reinder Vos De Wael, Konrad Wagstyl, et al. 2019. “Microstructural and Functional Gradients Are Increasingly Dissociated in Transmodal Cortices.” PLOS Biology 17 (5): e3000284. 10.1371/journal.pbio.3000284.

Park, Jinhan, Rachel L. M. Ho, Wei-en Wang, Shannon Y. Chiu, Young Seon Shin, and Stephen A. Coombes. 2025. “Age-Related Changes in Neural Oscillations Vary as a Function of Brain Region and Frequency Band.” Frontiers in Aging Neuroscience 17 (February). 10.3389/fnagi.2025.1488811.

Sala-Llonch, Roser, David Bartrés-Faz, and Carme Junqué. 2015. “Reorganization of Brain Networks in Aging: A Review of Functional Connectivity Studies.” Frontiers in Psychology 6 (May). 10.3389/fpsyg.2015.00663.

Sanz Leon, Paula, Stuart A. Knock, M. Marmaduke Woodman, et al. 2013. “The Virtual Brain: A Simulator of Primate Brain Network Dynamics.” Frontiers in Neuroinformatics 7 (June). 10.3389/fninf.2013.00010.

Schulz, Robert, Maximilian J. Wessel, Máximo Zimerman, Jan E. Timmermann, Christian Gerloff, and Friedhelm C. Hummel. 2015. “White Matter Integrity of Specific Dentato-Thalamo-Cortical Pathways Is Associated with Learning Gains in Precise Movement Timing.” Cerebral Cortex 25 (7): 1707–14. 10.1093/cercor/bht356.

Shafiei, Golia, Sylvain Baillet, and Bratislav Misic. 2022. “Human Electromagnetic and Haemodynamic Networks Systematically Converge in Unimodal Cortex and Diverge in Transmodal Cortex.” PLOS Biology 20 (8): e3001735. 10.1371/journal.pbio.3001735.

Shafiei, Golia, Ben D. Fulcher, Bradley Voytek, Theodore D. Satterthwaite, Sylvain Baillet, and Bratislav Misic. 2023. “Neurophysiological Signatures of Cortical Micro-Architecture.” Nature Communications 14 (1): 6000. 10.1038/s41467-023-41689-6.

Smith, Robert E., Jacques-Donald Tournier, Fernando Calamante, and Alan Connelly. 2015. “SIFT2: Enabling Dense Quantitative Assessment of Brain White Matter Connectivity Using Streamlines Tractography.” NeuroImage 119 (October): 338–51. 10.1016/j.neuroimage.2015.06.092.

Sydnor, Valerie J., Bart Larsen, Danielle S. Bassett, et al. 2021. “Neurodevelopment of the Association Cortices: Patterns, Mechanisms, and Implications for Psychopathology.” Neuron 109 (18): 2820–46. 10.1016/j.neuron.2021.06.016.

Tadel, François, Sylvain Baillet, John C. Mosher, Dimitrios Pantazis, and Richard M. Leahy. 2011. “Brainstorm: A User-Friendly Application for MEG/EEG Analysis.” Computational Intelligence and Neuroscience 2011 (1): 879716. 10.1155/2011/879716.

Taylor, Jason R., Nitin Williams, Rhodri Cusack, et al. 2017. “The Cambridge Centre for Ageing and Neuroscience (Cam-CAN) Data Repository: Structural and Functional MRI, MEG, and Cognitive Data from a Cross-Sectional Adult Lifespan Sample.” NeuroImage, Data Sharing Part II, vol. 144 (January): 262–69. 10.1016/j.neuroimage.2015.09.018.

Thomas Yeo, B. T., Fenna M. Krienen, Jorge Sepulcre, et al. 2011. “The Organization of the Human Cerebral Cortex Estimated by Intrinsic Functional Connectivity.” Journal of Neurophysiology 106 (3): 1125–65. 10.1152/jn.00338.2011.

Valk, Sofie L., Ting Xu, Daniel S. Margulies, et al. 2020. “Shaping Brain Structure: Genetic and Phylogenetic Axes of Macroscale Organization of Cortical Thickness.” Science Advances 6 (39): eabb3417. 10.1126/sciadv.abb3417.

Vidaurre, Diego, Laurence T. Hunt, Andrew J. Quinn, et al. 2018. “Spontaneous Cortical Activity Transiently Organises into Frequency Specific Phase-Coupling Networks.” Nature Communications 9 (1): 2987. 10.1038/s41467-018-05316-z.

Vos de Wael, Reinder, Oualid Benkarim, Casey Paquola, et al. 2020. “BrainSpace: A Toolbox for the Analysis of Macroscale Gradients in Neuroimaging and Connectomics Datasets.” Communications Biology 3 (1): 103. 10.1038/s42003-020-0794-7.

Voytek, Bradley, Mark A. Kramer, John Case, et al. 2015. “Age-Related Changes in 1/f Neural Electrophysiological Noise.” Articles. Journal of Neuroscience 35 (38): 13257–65. 10.1523/JNEUROSCI.2332-14.2015.

Wagstyl, Konrad, Stéphanie Larocque, Guillem Cucurull, et al. 2020. “BigBrain 3D Atlas of Cortical Layers: Cortical and Laminar Thickness Gradients Diverge in Sensory and Motor Cortices.” PLOS Biology 18 (4): e3000678. 10.1371/journal.pbio.3000678.

Wagstyl, Konrad, Claude Lepage, Sebastian Bludau, et al. 2018. “Mapping Cortical Laminar Structure in the 3D BigBrain.” Cerebral Cortex 28 (7): 2551–62. 10.1093/cercor/bhy074.

Wilson, Hugh R., and Jack D. Cowan. 1972. “Excitatory and Inhibitory Interactions in Localized Populations of Model Neurons.” Biophysical Journal 12 (1): 1–24. 10.1016/S0006-3495(72)86068-5.

Wu, Jinglong, Lihua Ma, Di Luo, et al. 2024. “Functional and Structural Gradients Reveal Atypical Hierarchical Organization of Parkinson’s Disease.” Human Brain Mapping 45 (4): e26647. 10.1002/hbm.26647.

Xia, Mingrui, Jin Liu, Andrea Mechelli, et al. 2022. “Connectome Gradient Dysfunction in Major Depression and Its Association with Gene Expression Profiles and Treatment Outcomes.” Molecular Psychiatry 27 (3): 1384–93. 10.1038/s41380-022-01519-5.

Xia, Yunman, Mingrui Xia, Jin Liu, et al. 2022. “Development of Functional Connectome Gradients during Childhood and Adolescence.” Science Bulletin 67 (10): 1049–61. 10.1016/j.scib.2022.01.002.

Yarkoni, Tal, Russell A. Poldrack, Thomas E. Nichols, David C. Van Essen, and Tor D. Wager. 2011. “Large-Scale Automated Synthesis of Human Functional Neuroimaging Data.” Nature Methods 8 (8): 665–70. 10.1038/nmeth.1635.

Zhang, Xi-Han, Kevin M. Anderson, Hao-Ming Dong, et al. 2025. “The Cell-Type Underpinnings of the Human Functional Cortical Connectome.” Nature Neuroscience 28 (1): 150–60. 10.1038/s41593-024-01812-2.

Zhang, Yu, and Zhe Sage Chen. 2025. “Harnessing Electroencephalography Connectomes for Cognitive and Clinical Neuroscience.” Nature Biomedical Engineering 9 (8): 1186–201. 10.1038/s41551-025-01442-4.

Zhou, Yingyao, Bin Zhou, Lars Pache, et al. 2019. “Metascape Provides a Biologist-Oriented Resource for the Analysis of Systems-Level Datasets.” Nature Communications 10 (1): 1523. 10.1038/s41467-019-09234-6.

